# Combined inhibition of mTOR and PIKKs exploits replicative and checkpoint vulnerabilities to induce death of PI3K-activated triple-negative breast cancer cells

**DOI:** 10.1101/700625

**Authors:** Sameer S. Chopra, Anne Jenney, Adam Palmer, Mario Niepel, Mirra Chung, Caitlin Mills, Sindhu Carmen Sivakumaren, Qingsong Liu, Jia-Yun Chen, Clarence Yapp, John M. Asara, Nathanael S. Gray, Peter K. Sorger

## SUMMARY

Frequent mutation of genes in the PI3K/AKT/mTOR signaling pathway in human cancers has stimulated large investments in therapeutic drugs but clinical successes have been rare. As a result, many cancers with high PI3K pathway activity such as triple-negative breast cancer (TNBC) are still treated primarily with conventional chemotherapy. By systematically analyzing responses of TNBC cells to a diverse collection of PI3K pathway inhibitors, we find that one drug, Torin2, is unusually effective because it inhibits both mTOR and PI3K-like kinases (PIKKs). In contrast to mTOR-selective inhibitors, Torin2 exploits dependencies on several kinases for progression of S-phase and for cell cycle checkpoints, thereby causing accumulation of single-stranded DNA and death by replication catastrophe or mitotic failure. Thus, Torin2 and its analogs represent a mechanistically distinct class of PI3K pathway inhibitors that are uniquely cytotoxic to TNBC cells. This insight could be translated therapeutically by further developing Torin2 analogs or combinations of existing mTOR and PIKK inhibitors.

## HIGHLIGHTS

- Torin2-like dual mTOR/PIKK inhibitors are cytotoxic to PI3K-activated TNBC cells
- Live-cell imaging shows that Torin2 exploits replicative and checkpoint vulnerabilities
- Combined inhibition of mTORC1/2 and Chk1 with selective drugs mimics Torin2
- Computational models show the importance of S-phase drug interactions for cytotoxicity
- The mechanism of action of Torin2 suggests new therapeutic opportunities for TNBC

## INTRODUCTION

Triple-negative breast cancers (TNBCs) are high-grade, invasive mammary ductal carcinomas with a poor prognosis (Foulkes et al., 2010). Defined at a molecular level by the absence of estrogen/progesterone receptors and a lack of *HER2*-amplification, TNBCs commonly have a “basal-like” gene expression signature similar to that of basal myoepithelial cells in the normal mammary duct (Perou et al., 2000). Whereas targeted therapy is standard of care for hormone receptor-positive and *HER2*-amplified breast cancers, TNBCs are primarily managed with chemotherapy (Bianchini et al., 2016). Relapsed disease and chemoresistance are common and thus new treatments for TNBC are needed.

TNBCs have the highest inferred PI3K/AKT/mTOR signaling activity of all breast cancer subtypes (Cancer Genome Atlas Network, 2012), providing a rationale for use of drugs targeting PI3K pathway kinases. In TNBC, PI3K pathway activation results not only from *PIK3CA*-activating mutations, but more commonly from recurrent mutation, deletion or silencing of the tumor suppressors *PTEN* and *INPP4B*, which encode lipid phosphatases that dephosphorylate the second messengers PIP_3_ and PIP_2_ to limit activation of AKT (Fedele et al., 2010; Gewinner et al., 2009; Marty et al., 2008). PI3K pathway kinases have been targeted with >40 clinical-grade small-molecule drugs, but clinical results in multiple tumor types have been disappointing (Janku et al., 2018). PI3K pathway activity is essential in non-transformed cells, limiting the therapeutic index of drugs targeting this pathway (Chia et al., 2015). Isoform-selective PI3K inhibitors were developed in part to overcome this problem, but these have relatively sporadic activity as monotherapy for solid tumors. For example, PI3K pathway activation in several *PTEN*-deficient cell lines is blocked by knockdown of PI3K p110β (*PIK3CB*) (Wee et al., 2008), but only a subset of a larger collection of *PTEN*-mutant cell lines is sensitive to a PI3K p110β inhibitor (Ni et al., 2012). The efficacy of isoform-selective drugs may be limited by insufficient pathway inhibition or by feedback mechanisms that cause pathway reactivation (Elkabets et al., 2013; Schwartz et al., 2015).

One approach to overcome such limitations is to combine PI3K pathway inhibitors with other drugs. A combination of the PI3K p110α (*PIK3CA*) inhibitor alpelisib and the selective estrogen receptor degrader fulvestrant was recently approved by the FDA for hormone receptor-positive breast cancers with *PIK3CA* mutations (André et al., 2019). The polypharmacology of PI3K pathway inhibitors provides a useful starting point to identify new strategies for combination therapy (Knight et al., 2010). Aside from selective inhibitors of PI3Kα, β, δ or γ, most PI3K pathway drugs inhibit several PI3K isoforms and/or mTORC1 and mTORC2. Some also inhibit PI3K-like kinases (PIKKs), which have structurally similar ATP binding sites. For example, the PI3K/mTOR inhibitor dactolisib (NVP-BEZ235) and the mTORC1/2 inhibitors Torin1 and Torin2 also inhibit ATR, ATM, and/or DNA-PK (Liu et al., 2010, 2013; Toledo et al., 2011). PIKK inhibition may be particularly relevant to TNBC given that common genetic changes such as *TP53* mutation and *MYC* or *CCNE1* amplification cause increased dependency on PIKKs for cell cycle checkpoint function and for countering oncogene-induced replication stress (Cancer Genome Atlas Network, 2012; Lin et al., 2017; Toledo et al., 2011; Zeman and Cimprich, 2014).

In this paper, we systematically analyzed responses of TNBC cells with PI3K pathway activation to a diverse collection of PI3K pathway inhibitors. We found that most drugs neither fully block proliferation nor cause cell death. In contrast, the pre-clinical compound Torin2 induced apoptosis in all TNBC cell lines tested. To determine which among the four or more high affinity targets of Torin2 is responsible, we used a chemical genetic approach involving dose-response studies in multiple cell lines, analysis of structural analogues and pharmacological reconstitution with mixtures of selective inhibitors. Phenotypes were determined using single-cell, time-course and pulse-labeling assays. These studies show that inhibition of both mTORC1/2 and PIKKs is required for the high activity of Torin2 in TNBC.

In contrast to other PI3K pathway drugs, Torin2 causes accumulation of single-stranded DNA (ssDNA) in S-phase cells, high levels of DNA damage signaling and death due to replication catastrophe or mitotic failure. In principle, blocking cells at G1/S by inhibiting mTORC1/2 should reduce cell killing in S phase or in mitosis caused by PIKK inhibition. However, computational modeling and experiments show that this is not true in TNBC cells because mTORC1/2 and PIKK activities are each required for progression of S phase. Thus, co-targeting the two classes of kinases in TNBC cells with a single drug exploits replicative and checkpoint vulnerabilities to cause death.

## RESULTS

### Torin2 produces strong cytotoxic and anti-proliferative effects in TNBC cells

Breast cancer cell lines are classified as basal or luminal based on gene expression (Neve et al., 2006). We performed western blotting on 46 cell lines and found that those with basal-like gene expression had uniformly low levels of PTEN and/or INPP4B (**Figure S1A, Table S1**); five such lines were selected for compound screening (HCC1806, Hs 578T, HCC38, BT-549 and HCC70). A sixth basal line (BT-20) with an activating *PIK3CA* mutation (H1047R) but normal levels of PTEN and INPP4B was also screened. All six lines lack hormone receptors and *HER2* amplification and are homozygous for *TP53* mutations (**Table S2**). We assembled a panel of 23 PI3K pathway inhibitors that included clinically approved drugs, investigational agents and tool compounds; the MEK inhibitor trametinib served as a MAPK pathway-specific comparator (**Table S3**). The panel covers drugs having diverse mechanisms of targeting the PI3K pathway (**Figure 1A**). Sub-confluent cultures were treated for 72h with each drug at nine doses and viable cells counted by microscopy at time t=0 and 72h (**Figure S1B**). Dose-response data were analyzed using growth rate inhibition (GR) metrics, which quantify drug sensitivity while correcting for differences in cell division times (Hafner et al., 2016): GR_50_ is analogous to IC_50_ and measures potency, while GR_max_ is analogous to E_max_ and measures efficacy (maximal drug effect; **Figure S1C**). A value of GR_max_ <0 indicates net cell loss, a value of zero represents no change in viable cell number, and a value >0 indicates net cell gain. By convention, GR_max_ = 1 in control cultures, which were treated with DMSO only.

**Figure 1.**
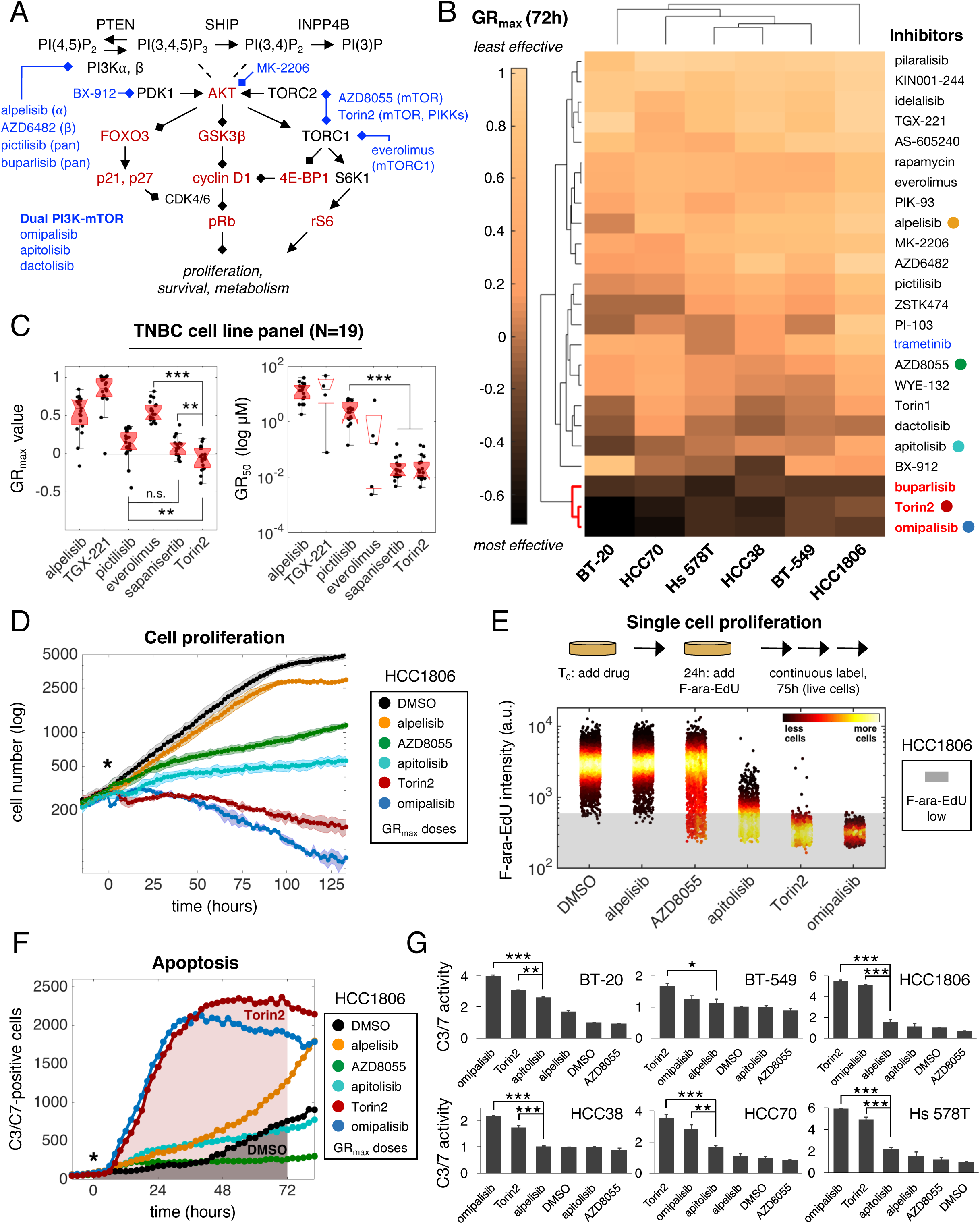
Torin2 produces strong cytotoxic and anti-proliferative effects in TNBC cells. **A**. Schematic of the PI3K pathway with select drugs (blue) and measured proteins (red). **B**. Clustergram heatmap of mean GR_max_ values for 24 drugs in six TNBC cell lines; N=3 experiments. Darker shading denotes GR_max_<0, which indicates cytotoxicity. Drugs in red are the most effective; the drug in blue is a MAPK inhibitor. Colored circles mark drugs analyzed in panels D-G. **C.** GR_max_ and GR_50_ values for the indicated drugs in a second analysis of 19 TNBC cell lines. *P<0.05, **P<0.01, ***P<0.001, and n.s. (not significant) by Mann-Whitney U test. **D**. Population growth curves for HCC1806 NLS-mCherry cells. “*” denotes start of treatment. Data points/shading depict mean ± SD of 3 replicates in a single time-lapse experiment; N=3 experiments, representative data shown. **E.** Mean nuclear F-ara-EdU intensity values in HCC1806 cells after treatment with various drugs for 24h followed by labeling for 75h; N=3 experiments, representative data shown. “a.u.” indicates arbitrary units. Shaded area indicates unlabeled cells. **F**. C3/C7-positive cell counts over time as detected by live-cell imaging of HCC1806 cells. Shading indicates AUC for 72h of drug exposure. **G**. C3/C7 activity (*i.e.*, mean fold-change in AUC ± SEM for drug vs. DMSO) for PI3K pathway drugs in six TNBC cell lines; N=2 experiments. *P<0.05, **P<0.01, ***P<0.001 by one-way ANOVA and Tukey’s test (only selected comparisons shown). Drugs were used at GR_max_ doses (1–3.2μM) in D, F and G. In E, Torin2 and omipalisib were used at 0.3xGR_max_ (1μM) due to cell death at higher doses.

Highly effective responses (negative GR_max_ values) were rare across the 144 drug-cell line combinations examined (**Figure 1B**), but three drugs were broadly cytotoxic: the phase 3 PI3K p110 pan-isoform inhibitor buparlisib, the phase 1 dual PI3K/mTOR inhibitor omipalisib and the pre-clinical dual mTOR/PIKK inhibitor Torin2. Omipalisib and Torin2, but not buparlisib, were also highly potent, with nanomolar GR_50_ values (**Figure S1D**). Torin2 is the least studied molecule of the three, but we found it to be more cytotoxic in 19 basal-like cell lines than sapanisertib, an mTORC1/2 inhibitor presently in clinical trials (**Figure 1C**). Torin2 exhibited similar efficacy to sapanisertib in luminal breast cancer cell lines (N=7), and caused little or no cytotoxicity in non-malignant mammary epithelial cells (N=2) (**Figure S1E**). Thus, Torin2 may represent an improved way to target TNBC.

To identify factors influencing drug response, we assayed proliferation and apoptosis in live cells. To enable a principled comparison, compounds were used at the lowest concentration eliciting a GR_max_ response (*i.e.*, the ‘GR_max_ dose’). We imaged HCC1806 cells expressing NLS-mCherry continuously for >5 days (∼4.5 cell divisions; division time T_d_=28h) in the presence of DMSO or several drugs whose GR_max_ values span a broad range. Drugs with positive GR_max_ values, such as alpelisib (GR_max_=0.92), AZD8055 (GR_max_=0.56) and apitolisib (GR_max_=0.37), reduced viable cell number but did not fully block proliferation (**Figure 1D**). Conversely, drugs with negative GR_max_ values such as Torin2 (GR_max_=-0.16) and omipalisib (GR_max_=-0.29) steadily reduced cell number. At a 10-fold lower dose (0.32µM) of Torin2 or omipalisib, there was still no increase in cell number over time, consistent with complete cytostasis or a balance between cell division and death (**Figure S1F**).

As another measure of proliferation, we exposed HCC1806 cells to drugs for 24h and then added F-ara-EdU, a non-toxic thymidine analog that is incorporated into newly-synthesized DNA, for an additional 75h (2.7xT_d_) (Neef and Luedtke, 2011). GR_max_ doses of alpelisib, AZD8055, and apitolisib were only partly effective at inhibiting labeling during this period, resulting in 0.7%, 6.3%, and 65.4% F-ara-EdU-negative cells, respectively; in contrast, Torin2 and omipalisib each inhibited labeling of >99% of viable cells (**Figures 1E, S1G**). Thus, the low GR_max_ values of Torin2 and omipalisib are associated with a marked reduction in the number of viable cells that actively synthesize DNA at any time during an extended period of drug exposure.

To assay induction of apoptosis, we imaged all six selected cell lines following drug exposure in the presence of a fluorogenic substrate of executioner caspases 3 and 7 (C3/C7) (**Figure S1H**). We then calculated the fold-change in the area under the fluorescence curve (AUC) between 0 and 72h relative to a DMSO control (**Figures 1F-G**). Drugs such as AZD8055 induced no detectable C3/C7 activity in any cell line. In contrast, Torin2 and omipalisib increased C3/C7 activity by an average of ∼3.5-fold across all six cell lines. In summary, the data show that low GR_max_ values for Torin2 and omipalisib in TNBC cells result from effective inhibition of proliferation and induction of apoptosis.

### PI3K/AKT/mTOR inhibitors impede progression of S phase

To evaluate the effects of Torin2 on signaling, we exposed TNBC cells to GR_max_ doses of drugs for 24h and then quantified the levels or localization of nine proteins/phosphoproteins by immunofluorescence microscopy (**Figures 1A, 2A, S2A-C**). AKT activity was inferred from (i) phosphorylation of AKT at T308 and S473, (ii) phosphorylation of the AKT substrate GSK3β at S9 and (iii) nuclear localization of FOXO3, which is regulated by AKT. AKT^473^ phosphorylation served as a measure of mTORC2 activity, 4E-BP1^T37/T46^ as a measure of mTORC1 activity and ribosomal protein S6^S235/S236^ phosphorylation as a measure of S6K1 activity. We also measured levels of cyclin D1 and the CDK inhibitors p21 and p27, which are regulated by the PI3K pathway. Based on the magnitude of changes in the levels of these proteins/phosphoproteins, Torin2 and omipalisib were the most effective drugs at inhibiting the PI3K pathway (**Figures 2A, S2D**). To assay the net “output” of the PI3K pathway, we quantified CDK-dependent phosphorylation of the retinoblastoma protein on S807/811 (p-pRb). When mitogenic signaling by PI3K and/or other pathways is low, CDK activity is also low, pRb is dephosphorylated and E2F is inhibited, causing G1 arrest (Duronio and Xiong, 2013). In HCC1806 cells, exposure to Torin2 or omipalisib for t=T_d_ increased the percentage of p-pRb-low cells from 5% to 64% and 81%, respectively, compared to 6-42% for six other PI3K pathway drugs tested (**Figures 2B, S2E**). Thus, low GR_max_ values for Torin2 and omipalisib correlate with greater suppression of PI3K signaling and lower levels of p-pRb (Liang and Slingerland, 2003).

**Figure 2.**
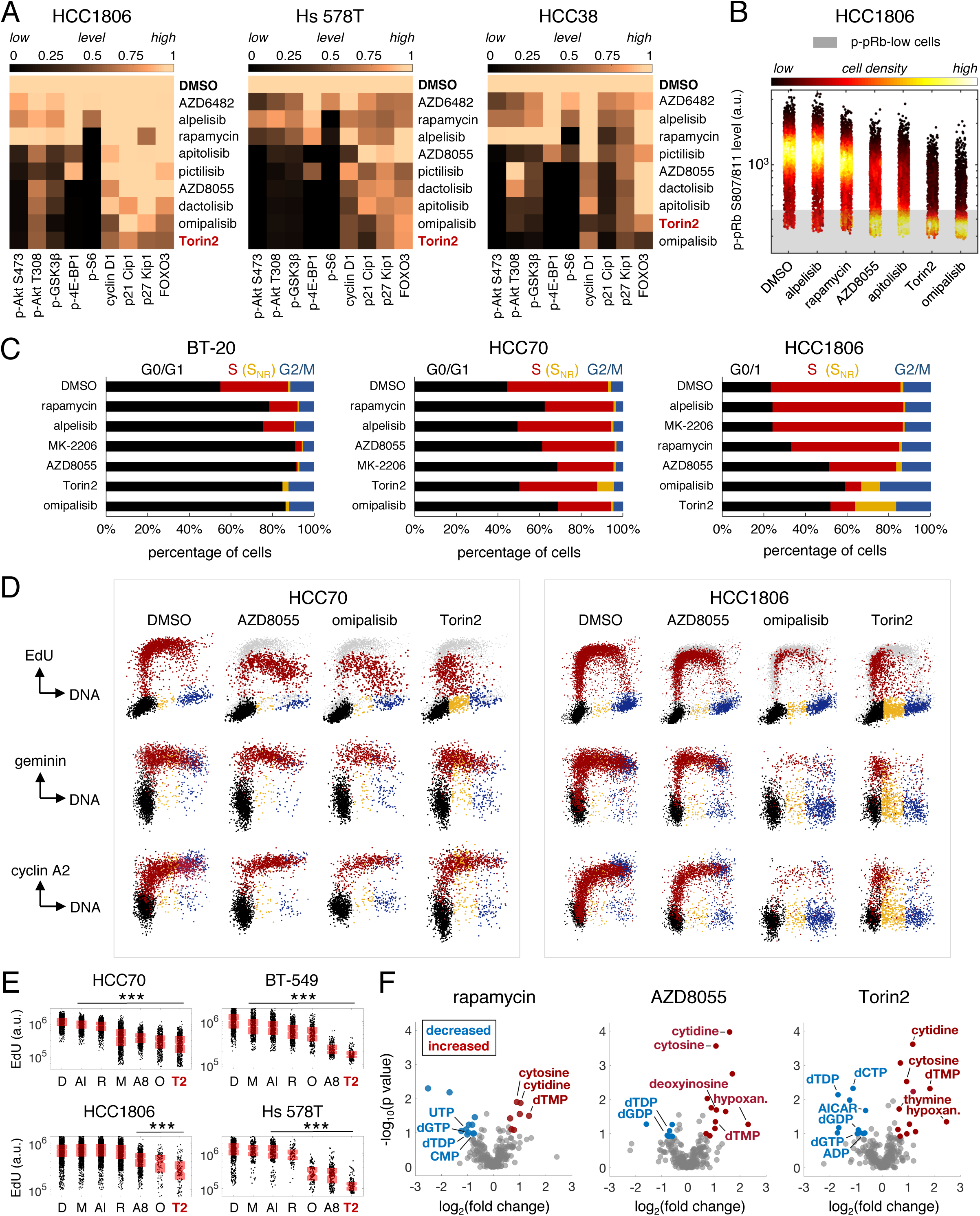
PI3K/AKT/mTOR inhibitors impede progression of S phase. **A**. Clustergram heatmaps rank the activity of PI3K pathway drugs based on mean values of the levels and/or localization of nine proteins/phosphoproteins at 24h post-treatment. Data are normalized to DMSO; N≥2 experiments. Drugs were used at GR_max_ doses (1-3.2μM) except for Torin2 and omipalisib, which were used at 0.3xGR_max_ (1μM) in HCC1806 cells due to cell death. **B.** Mean nuclear intensity values of phospho-pRb S807/811 (p-pRb) in HCC1806 cells at t=T_d_ (28h) post-treatment. **C**. Cell cycle distributions at t=T_d_ post-treatment for BT-20 (48h), HCC70 (45h) and HCC1806 (28h) based on analysis of nuclear DNA content vs EdU content. Colors indicate cell cycle stage: G1 (black), active S phase (red), S-phase non-replicating (S_NR_; yellow), and G2/M (blue). Drugs are ordered from least effective to most effective (top to bottom) based on viable cell counts at t=T_d_. GR_max_ doses (1–3.2μM) were used for all drugs except Torin2 and omipalisib, which were used at 0.3xGR_max_ dose (BT-20: 1μM; HCC70: 0.32–1μM; HCC1806: 1μM) due to cell death. **D**. *Top row*: Nuclear DNA content vs EdU content in HCC70 and HCC1806 cells at t=T_d_. Data points for DMSO (grey) are shown in background for comparison. Doses were as noted in C. *Middle, bottom rows*: Nuclear DNA content vs. mean nuclear intensity values for geminin or cyclin A2 at t=T_d_. Colors indicate different cell cycle stages as determined by DNA content and EdU content values measured in the same cells. In HCC70 only, Torin2 and omipalisib were used at 0.1xGR_max_ dose (0.1–0.32μM) due to cell death. **E**. Quantification of total nuclear EdU content at time=T_d_ in gated S-phase cells (red cells, panel D, top row; S_NR_ cells excluded). For HCC70 and HCC1806 cells, doses were as noted in C. For Hs 578T cells, GR_max_ doses (1–3.2μM) were used for all except Torin2 and omipalisib, which were used at 0.3xGR_max_ dose (1μM). For BT-549 cells, AZD8055 was used at 0.3xGR_max_ dose (0.32μM), and omipalisib and Torin2 were used at 0.03xGR_max_ dose (0.1μM) due to low numbers of S-phase cells at higher doses. Al:alpelisib, A8:AZD8055, D:DMSO, M:MK-2206, O:omipalisib, R:rapamycin, T2:Torin2. ***, P<0.001 for each drug vs DMSO (Mann-Whitney U test). **F**. Volcano plots of log_2_(fold change) vs. –log_10_(p-value) for intracellular polar metabolites after exposure of BT-549 cells to GR_max_ doses of rapamycin or AZD8055 or 0.3xGR_max_ dose of Torin2 (1μM for all three drugs) for t=0.15xT_d_ (6h). Downregulated metabolites appear in blue, upregulated metabolites appear in red. Nucleotides and precursors are labeled.

To determine if Torin2 inhibits cell cycle progression at G1/S to a greater extent than other PI3K pathway drugs, we performed two-parameter cell cycle analysis by staining DNA with Hoechst and pulse labeling S-phase cells with EdU (Figure S3A). Cell lines with different division times were compared by exposing cells to drugs for t=T_d_ and then pulse-labeling with EdU for 0.025xT_d_ (40-70min). EdU-positive cells were scored as being in S phase and EdU-negative cells scored as being in G0/G1 or G2/M based on DNA content. EdU-negative cells with DNA content intermediate between G0/G1 and G2/M cells were scored as S-phase-non-replicating (S_NR_) cells (Shi et al., 2001).

Exposure of BT-20 cells to Torin2 or omipalisib at 0.3xGR_max_ dose (1µM) for t=T_d_ (48h) increased the G1 fraction to ∼85% (from 55% in DMSO control) and reduced the S-phase fraction to 0% (from 33% in DMSO control; **Figure 2C**). In contrast, GR_max_ doses of other drugs that were less effective at suppressing PI3K pathway activity (such as alpelisib and rapamycin) increased the G0/G1 fraction to a lesser degree (76–79%) and incompletely reduced the S-phase fraction (to 13–15%). Thus, in *PIK3CA*-mutant TNBC cells, the high activity of Torin2 and omipalisib is associated with an increased blockade of cells at G1/S and elimination of S-phase cells.

In contrast, exposure of other TNBC cell lines to Torin2 or omipalisib, even at doses that blocked proliferation, did not eliminate S-phase cells (**Figures 2C, S3B**). For example, 26-41% of HCC70 cells were in S phase after exposure to Torin2 or omipalisib for t=T_d_ (45h). To determine if this phenotype arises because Torin2 and other PI3K pathway drugs impede progression of cells through S phase, we quantified the amount of EdU incorporated into newly-synthesized DNA during pulse-labeling. To identify S-phase cells independent of DNA synthesis, we stained for expression of the APC/C substrates geminin and cyclin A2 by immunofluorescence. In HCC70 cells, we found that multiple PI3K pathway drugs reduced EdU content in geminin- and cyclin A2-positive cells. Torin2 suppressed DNA synthesis to the greatest degree and caused accumulation of S_NR_ cells (∼9% of all cells; **Figures 2D-E**). In HCC1806 cells, Torin2 elicited even stronger inhibition of S phase in a dose-dependent manner, with peak accumulation of 20% S_NR_ cells (**Figures 2D-E, S3C-D**). These S_NR_ cells exhibited decreased levels of geminin, cyclin A2 and p-pRb, consistent with cell cycle exit (**Figures 2D, S3E**). PI3K pathway drugs also reduced total EdU content in BT-549 and Hs 578T cells, with Torin2 again having the strongest effect (**Figures 2E, S3F-G**). The same was true of BT-20 cells exposed to drugs for less time (t=0.125-0.3xT_d_; 6-14h), well in advance of the G0/G1 arrest that occurred by t=T_d_ (**Figures S3H-I**). Thus, Torin2 and other PI3K pathway drugs not only inhibit the cell cycle at G1/S, but also interfere with progression of cells through S phase.

The PI3K pathway is known to play a role in nucleotide metabolism (Wang et al., 2009), and we hypothesized that exposure of TNBC cells to PI3K pathway drugs inhibits S phase by decreasing levels of DNA precursors (Juvekar et al., 2016a). To test this idea, we performed targeted metabolomic profiling by mass spectrometry following a short exposure of BT-549 cells to one of three mTOR inhibitors (rapamycin, AZD8055, or Torin2) or DMSO for t=0.15xT_d_ (6h; **Figure 2F, Table S4**). Drug-exposed cells exhibited changes in the levels of multiple purine and pyrimidine deoxyribonucleotides and their precursors, consistent with inhibition of both *de novo* synthesis and salvage pathways. Torin2 elicited the largest number of such changes, but all three mTOR inhibitors caused increased levels of cytidine and cytosine, key substrates in the salvage of pyrimidines, as well as increased levels of deoxythymidine monophosphate (dTMP) and decreased levels of deoxythymidine diphosphate (dTDP), consistent with reduced activity of deoxythymidylate kinase, an enzyme that acts downstream of both *de novo* pyrimidine synthesis and salvage pathways. Thus, it is likely that Torin2 and other drugs tested inhibit S-phase progression at least in part by depleting TNBC cells of nucleotides required for DNA replication.

### Torin2 causes substantial cell killing during S/G2

To better understand how Torin2 acts on cells in different cell cycle stages, we performed time-lapse imaging of asynchronous HCC1806 cells stably expressing H2B-mTurquoise and mVenus-hGeminin(1-110), a reporter of cell cycle stage (Sakaue-Sawano et al., 2008). Cells were exposed to DMSO or 0.3xGR_max_ dose (1µM) of omipalisib or Torin2 and then followed for 48h (N=340 single cells tracked and analyzed; **Figure 3A**). H2B-mTurquoise was used to identify nuclei, monitor cell division and score cell death by nuclear fragmentation. Cell cycle stage was determined by the fluorescence intensity level of mVenus, which is low in G0/G1 and high in S/G2. In the presence of DMSO, single cells divided every 24.2±0.8h, consistent with population doubling times, and only ∼6% died (**Figure 3A**, grey bars). Exposure to omipalisib caused death of 38% of cells (N=43/112) and blocked progression of 55% (N=61/112); the remainder of cells were incompletely blocked (**Figures 3A-B**). Torin2 was more cytotoxic, killing 51% of cells (N=62/122) and blocking progression of 43% (N=53/122).

**Figure 3:**
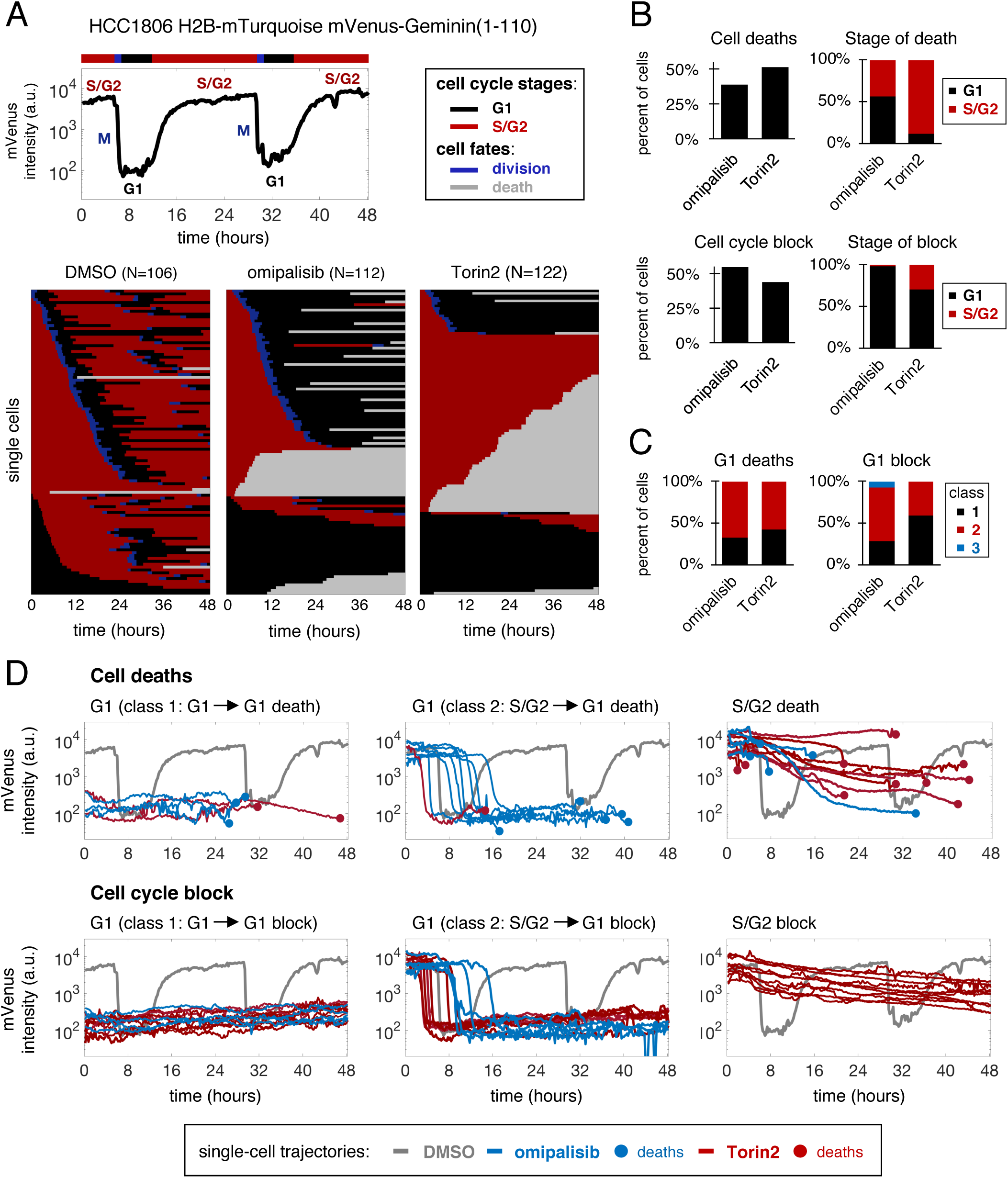
Torin2 causes substantial cell killing during S/G2. **A.** Time-lapse imaging of asynchronous HCC1806 cells expressing H2B:mTurquoise and mVenus:hGeminin(1-110). mTurquoise was used to score cell division and death; mVenus intensity levels were used to identify cell cycle stage, as illustrated for a representative cell exposed to DMSO. Heatmaps show the progression of single cells through the cell cycle after exposure to DMSO or a 0.3xGR_max_ dose (1μM) of omipalisib or Torin2 for 48h. Colors denote cell cycle stages and cell fates, as shown in the figure legend. **B**. Left plots show the frequency of deaths and cell cycle block caused by omipalisib and Torin2; right plots show the frequency of these events by cell cycle stage. **C**. Frequencies of different classes of responses leading to G1 death or block. *Class 1*: a cell in G1 at the start of treatment is killed or blocked in the same G1; *class 2*: a cell in S/G2 at start of treatment proceeds to division, and the tracked daughter cell is killed or blocked in the following G1; *class 3*: similar to class 2, except the cell starts in G1. **D**. Phenotypic responses caused by drug exposure. Shown are representative traces of mVenus fluorescence intensity vs. time for single cells (omipalisib in blue, Torin2 in red). For comparison, the trace for a representative cell exposed to DMSO is shown in the background (grey).

Confirming data on fixed cells, live-cell imaging showed that Torin2 and omipalisib are active during both G1 and S/G2 phases of the cell cycle. In the presence of omipalisib, similar numbers of cells died in S/G2 (44%; N=19/43) as in G1 (56% of cells; N=24/43). In contrast, 8-fold more cells exposed to Torin2 died in S/G2 (89% of cells, N=55/62) than in G1 (11%, N=7/62) (**Figures 3A-B**). Among surviving cells, Torin2 also imposed a stronger S/G2 block than omipalisib: 30% of blocked cells (N=16/53) were in S/G2 after exposure to Torin2 compared to only 2% (N=1/61) for omipalisib. The strong effects of Torin2 on S/G2 cells caused the sequence of events leading to G1 block or death to differ between the two drugs. For example, the majority of omipalisib-treated cells that died in G1 (66.7%, N=16/24) or remained blocked there (65%, N=39/60) were in S/G2 at the time of drug exposure; they then proceeded through mitosis to the following G1 where they either died or arrested (G1 “class 2” death or block; **Figures 3C-D**). In contrast, G1 class 2 deaths (57.1%, N=4/7) and block (40.5%, N=15/37) were less frequent in the presence of Torin2 because the majority of cells in S/G2 at the time of initial drug exposure never reached mitosis. We conclude that both Torin2 and omipalisib are active on cells in G1 and S/G2, but that exposure to Torin2 causes substantially greater arrest and death of S/G2 cells than omipalisib.

### Torin2 causes replication catastrophe

Impaired progression of S phase by nucleotide insufficiency or other genotoxic insults is a cardinal manifestation of “replication stress” (Zeman and Cimprich, 2014). Stalled replication forks cause accumulation of ssDNA because the activities of DNA polymerases and replicative helicases become uncoupled; excess ssDNA can then consume replication factors and cause DNA breakage and death by “replication catastrophe” (Toledo et al., 2013; Zeman and Cimprich, 2014). To determine if exposure to Torin2 causes accumulation of ssDNA, HCC1806 cells were labeled with BrdU for t=T_d_, treated with GR_max_ doses of drug for t=0.2xT_d_ (5h) and immunostained with anti-BrdU antibodies under non-denaturing (“native”) conditions. Exposure to Torin2 – but not to omipalisib or AZD8055 – increased the fraction of cells with elevated ssDNA ∼8-fold (**Figures 4A**, labeled red; **4C**) and mean nuclear BrdU intensity in gated S-phase cells ∼2-fold (**Figure 4B**). Similar increases were observed with AZ20 and rabusertib, selective inhibitors of ATR kinase and its substrate kinase Chk1, which act together to stabilize stalled replication forks and suppress origin firing (Zeman and Cimprich, 2014). Exposure of HCC1806 cells to Torin2, AZ20 and rabusertib also increased levels of phosphorylated H2A.X^S139^ (γH2A.X), a marker of DNA damage, and of phosphorylated RPA^S4/8^ (p-RPA), an ssDNA-binding protein (**Figures 4A-C**). Similar effects were seen in HCC70 cells but after longer periods of drug exposure (∼0.5xT_d_, 24h) (**Figures S4A-B**). In HCC1806 cells, these responses were associated with activated intra-S-phase checkpoint signaling, as evidenced by a ∼15-fold increase in the fraction of cells with elevated nuclear p-Chk1^S317^ and p-Chk2^T68^ (**Figures 4C, S4C**). In contrast, neither AZD8055 nor omipalisib caused any detectable increase in ssDNA, γH2A.X, p-RPA, p-Chk1^S317^ or p-Chk2^T68^, despite inhibiting DNA synthesis (**Figure 2**). Thus, Torin2 is unique among the PI3K pathway drugs we tested in causing accumulation of ssDNA, DNA damage and increased checkpoint signaling in TNBC cells.

**Figure 4:**
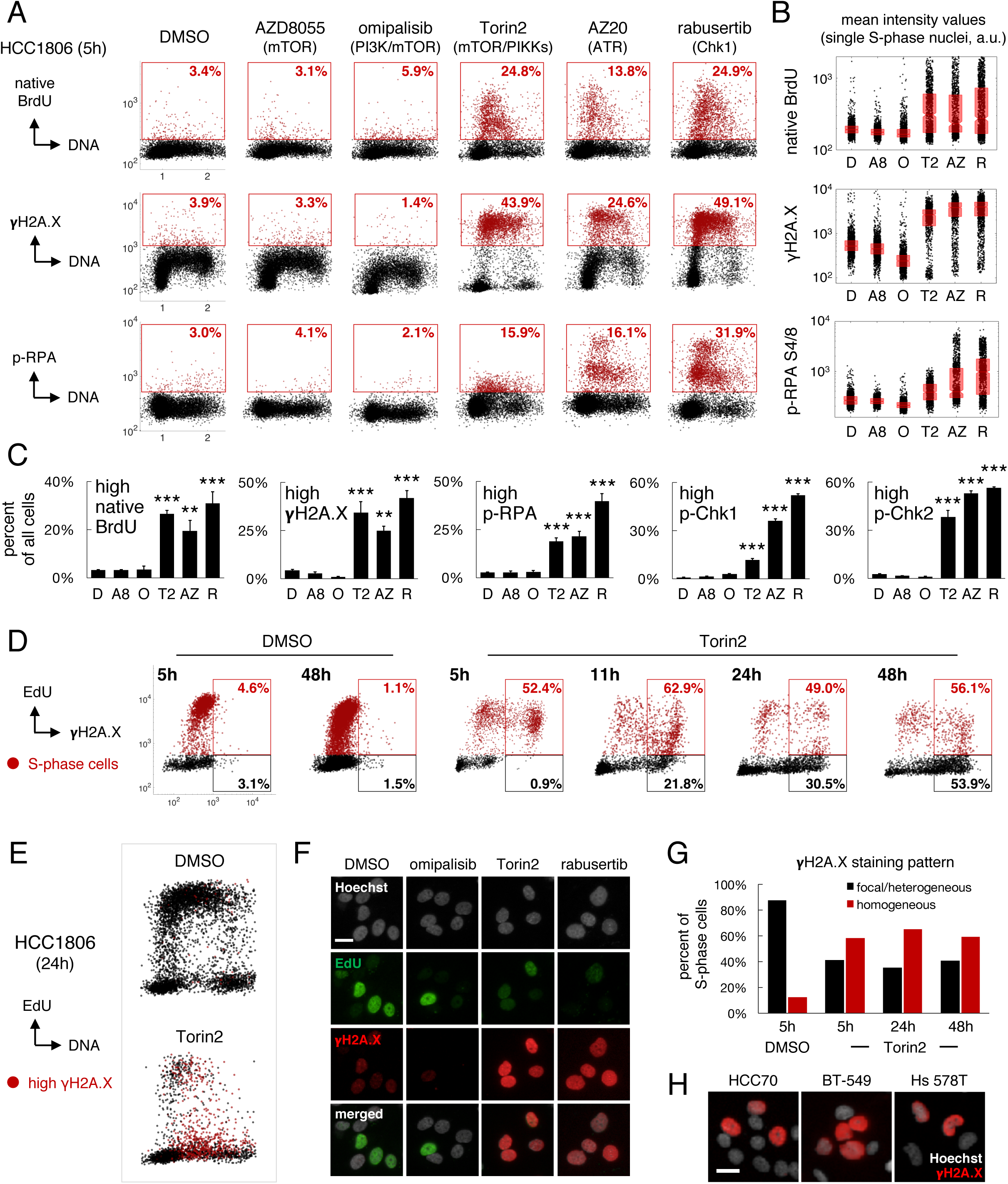
Torin2 causes replication catastrophe. HCC1806 cells were exposed to 0.3-1xGR_max_ drug doses (1–3.2μM) throughout. **A.** DNA content vs. intensities of native BrdU, γH2A.X or p-RPA S4/8 after exposure to the indicated drugs for ∼0.2xT_d_ (5h). Cells with increased levels are gated and marked in red. Percentages are of the entire treated population. **B**. Quantification of nuclear intensity values of native BrdU, γH2A.X, and p-RPA in gated S-phase cells at 5h post-treatment. Boxplots show median and 25th/75th percentiles. D:DMSO, A8:AZD8055, O:omipalisib, T2:Torin2, AZ:AZ20, R:rabusertib. **C**. Mean percent ± SEM of cells with increased levels of native BrdU, γH2A.X, p-RPA S4/8, p-Chk1 S317 and p-Chk2 T68; N=3 experiments. **P<0.01, ***P<0.001 vs DMSO by one-way ANOVA and Dunnett’s test. **D**. Mean intensity for EdU vs. γH2A.X after treatment with DMSO or 1μM Torin2 for the indicated times. EdU/γH2A.X double-positive cells (red gate) and EdU-negative/γH2A.X-positive cells (black gate) are quantified as a percent of all EdU-positive (red) or EdU-negative (black) cells, respectively. **E**. DNA content vs mean EdU intensity at 24h post-treatment with DMSO or 1μM Torin2. Cells with elevated γH2A.X levels (> 10^3^ a.u.) are marked in red. **F**. Nuclear γH2A.X staining pattern in S-phase/S_NR_ cells at 24h post-treatment with drugs at 1μM. Pan-nuclear γH2A.X staining is an indicator of high genotoxic stress. **G**. Percent of S-phase cells with pan-nuclear γH2A.X staining (red) vs. more focal staining (black) after exposure to DMSO or 1μM Torin2. **H.** γH2A.X staining pattern in various TNBC cell lines after treatment with 1μM Torin2. Scale bars in F and H=20μm.

To determine whether increased ssDNA and DNA damage precede accumulation of S_NR_ cells and death, HCC1806 cells were exposed to Torin2 at 0.3xGR_max_ dose for 0.2–1.7xT_d_ (5–48h). At 5, 11, 24 or 48h, cells were pulse-labeled with EdU for 0.025xT_d_ (40min) and counterstained with anti-γH2A.X antibodies. Between 5 and 48h of drug exposure, ∼50–60% of EdU-positive cells were positive for γH2A.X, representing a greater than 10–fold increase over cells exposed to DMSO for 5h (**Figure 4D**). Over the same time period, the fraction of EdU-negative cells that were γH2A.X-positive increased from 1% to 54%; these cells had DNA content values characteristic of S_NR_ cells (**Figure 4E**). Similar to rabusertib, Torin2 exposure resulted in S_NR_ cells exhibiting a homogeneous, pan-nuclear γH2A.X staining pattern associated with severe genotoxic stress and commitment to apoptosis (de Feraudy et al., 2010) (**Figure 4F**). No such staining pattern was observed after cells were exposed to omipalisib. More than 50% of S-phase cells exposed to Torin2 for 5–48h exhibited a pan-nuclear γH2A.X staining pattern (**Figure 4G**). Similar effects were observed in other TNBC cell lines (**Figure 4H**). Thus, ssDNA and DNA damage induced by Torin2 precede the appearance of S_NR_ cells exhibiting hallmarks of replication catastrophe.

### The activity of Torin2 results from combined inhibition of mTORC1/2 and PIKKs

To test the hypothesis that the activity of Torin2 on TNBC cells is a consequence of its unique polypharmacology (Liu et al., 2013), we attempted to reconstitute its effects in HCC1806 and HCC70 cells by combining selective inhibitors of mTOR and PIKKs. The combination of AZD8055 and rabusertib more closely matched the activity of Torin2 across a range of doses than AZD8055 and AZ20 (**Figures 5A, S5A**), possibly because inhibition of Chk1 mimics the inhibition of multiple PIKKs (Buisson et al., 2015). Nonetheless, AZD8055 combined with either rabusertib or AZ20 was more effective (by GR_max_) than inhibiting mTORC1/2 alone and more potent (by GR_50_) than inhibiting PIKKs alone. In a complementary approach, we compared AZD8055 to nine Torin2 analogs (**Figure S5B**) and to Torin1, which inhibits mTORC1/2 and DNA-PK but not ATR or ATM (Liu et al., 2010) (**Figures 5B-C**). Most Torin2 analogs were less potent than AZD8055 and Torin1 in HCC1806 and HCC70 cells, but more effective (*i.e.*, negative GR_max_ values).

**Figure 5:**
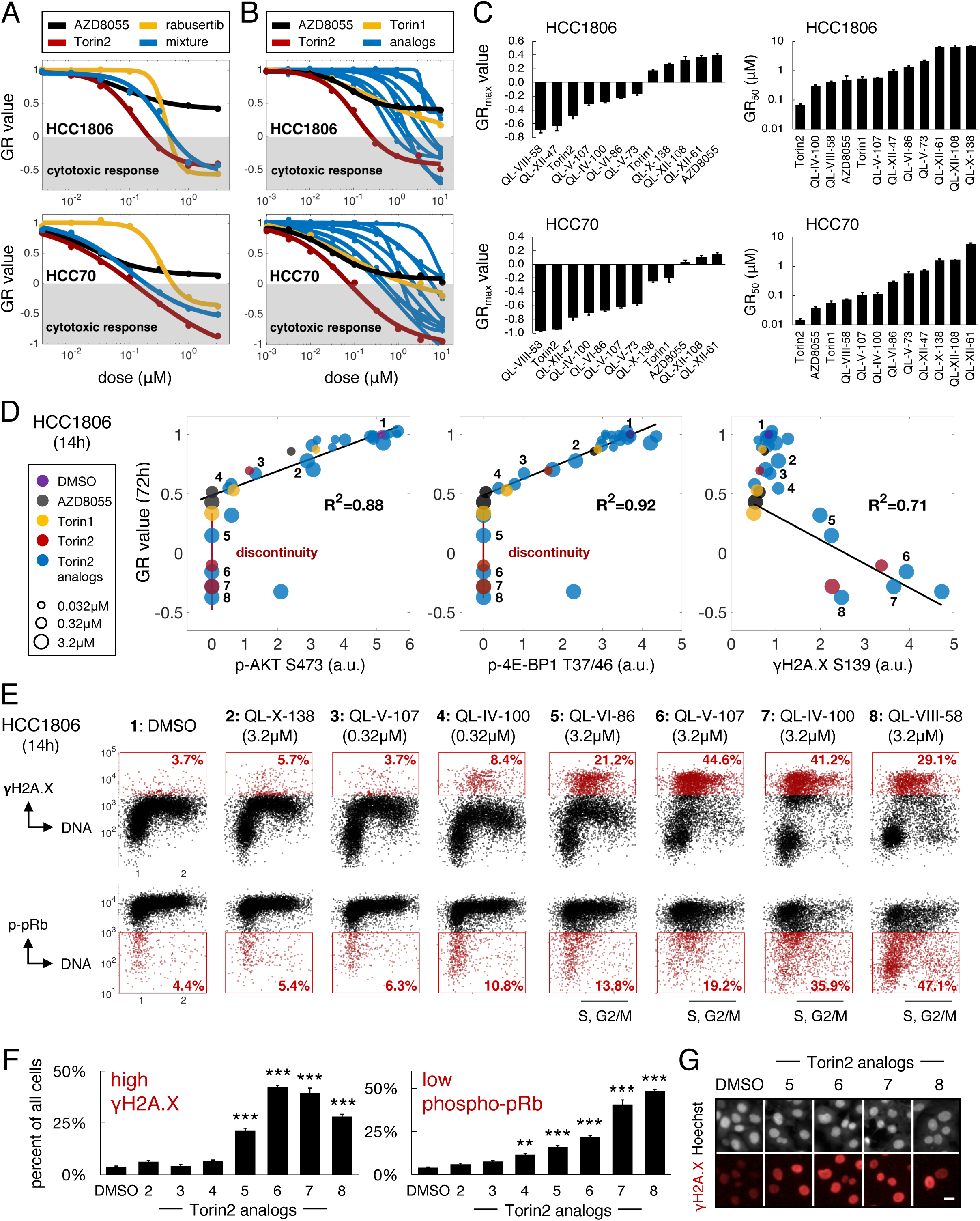
The activity of Torin2 results from combined antagonism of mTORC1/2 and PIKKs. **A.** Dose-response curves (72h) for AZD8055, Torin2, rabusertib, and AZD8055/rabusertib mixed at equimolar concentrations in HCC1806 and HCC70 cells. **B**. Dose-response curves for AZD8055, Torin1, Torin2 and nine different Torin2 analogs (72h). **C**. Mean ± SEM of GR_max_ and GR_50_ values for Torin1, Torin2, AZD8055 and Torin2 analogs in HCC1806 and HCC70 cells; N=3 experiments. **D**. Levels of p-AKT S473, p-4E-BP1 T37/46 and γH2A.X at t=0.5xT_d_ (14h) post-treatment vs. GR values computed after 72h of exposure of HCC1806 cells to 0.032–3.2μM of AZD8055, Torin1, Torin2 or Torin2 analogs. Drugs/doses whose GR values correlate with mTORC1/2 signaling activity alone fall on the black regression lines (*left, middle*). Drugs/doses whose activities are discontinuous with the black regression line fall on the vertical red lines (*left, middle*) and cause increased levels of γH2A.X (*right*). **E**. DNA content vs. nuclear γH2A.X or p-pRb levels at 14h for DMSO (#1) and drugs/doses whose activities correlate with mTORC1/2 signaling activity (#2-4) and those whose activities do not (#5-8). Percentages of gated γH2A.X-high or p-pRb-low cells are relative to entire treated population. **F**. Mean percent ± SEM of γH2A.X-high cells or p-pRb-low cells at 14h post-treatment with various Torin2 analogues; N=3 experiments. **P<0.01, ***P<0.001 by one-way ANOVA and Dunnett’s test. **G**. Nuclear γH2A.X staining pattern in S-phase cells after exposure to indicated treatments for 14h. Scale bar=20μm.

To dissect the relative contribution of mTORC1/2 and PIKK inhibition to the activity of Torin2 analogs, we exposed HCC1806 and HCC70 cells to a 100-fold concentration range of each drug for t=0.33–0.5xT_d_ (14h) and assayed activity against mTORC1/2 by immunofluorescence for p-AKT^S473^ and p-4E-BP1^T37/T46^. We found that the magnitude of mTORC1/2 inhibition at 14h correlated well with GR values at 72h for multiple drugs and analogs, including Torin1 and AZD8055 (**Figures 5D, S5C,** black regression lines). However, at low levels of mTORC1/2 signaling activity, the GR values for Torin2 (red circles) and the analogs QL-VI-86 (#5), QL-V-107 (#6), QL-IV-100 (#7) and QL-VIII-58 (#8) were substantially lower, resulting in a discontinuous relationship between mTORC1/2 inhibition and GR values (**Figures 5D, S5C**, red lines/boxes). Thus, mTORC1/2 inhibition alone does not fully account for the high activity of these drugs. This same subset of drugs caused elevated pan-nuclear staining of γH2A.X in S-phase cells, consistent with inhibition of PIKKs (**Figures 5D-G, S5C-E**). In HCC1806 cells, these drugs also produced increased fractions of p-pRb-low cells in G1, S and G2/M phases, indicative of arrest at multiple points in the cell cycle (**Figures 5E**, see black line beneath individual scatterplots, and **5F**). We conclude that Torin2 and its analogs exhibit more or less of two distinct cellular activities. Whereas all compounds inhibit mTORC1/2 and cause growth inhibition, a subset of compounds including Torin2 and #5–8 also strongly inhibit PIKKs to cause DNA damage and cytotoxicity. The most effective drugs are those with the strongest combined activity.

### Combined inhibition of mTORC1/2 and PIKKs produces benefit by co-targeting pathways required in S phase

The high activity of Torin2 and related dual mTOR/PIKK inhibitors is unexpected. In principle, G0/G1 arrest caused by mTORC1/2 inhibition should antagonize cell death in S phase from PIKK inhibition (Johnson et al., 1999; King et al., 2015). To better understand why Torin2 and its analogs are so effective, we developed two cell cycle models based on discrete time-step *Monte Carlo* simulations. In these models, single cells progressed through G1, S, G2 and M “compartments” at rates matching experimental data. Following M phase, each mother cell was replaced by two G1 daughters. In the “*independence model*,” mTORC1/2 inhibition slowed transit only through G1 (**Figure 6A**), while in the “*interaction model,*” mTORC1/2 inhibition slowed transit through both G1 and S phase (**Figure 6B**). In both models, ATR/Chk1 inhibition caused lethal replication block. GR values and total cell killing (“cell loss factor”) were then calculated for simulations with different degrees of mTORC1/2 and ATR/Chk1 inhibition.

**Figure 6:**
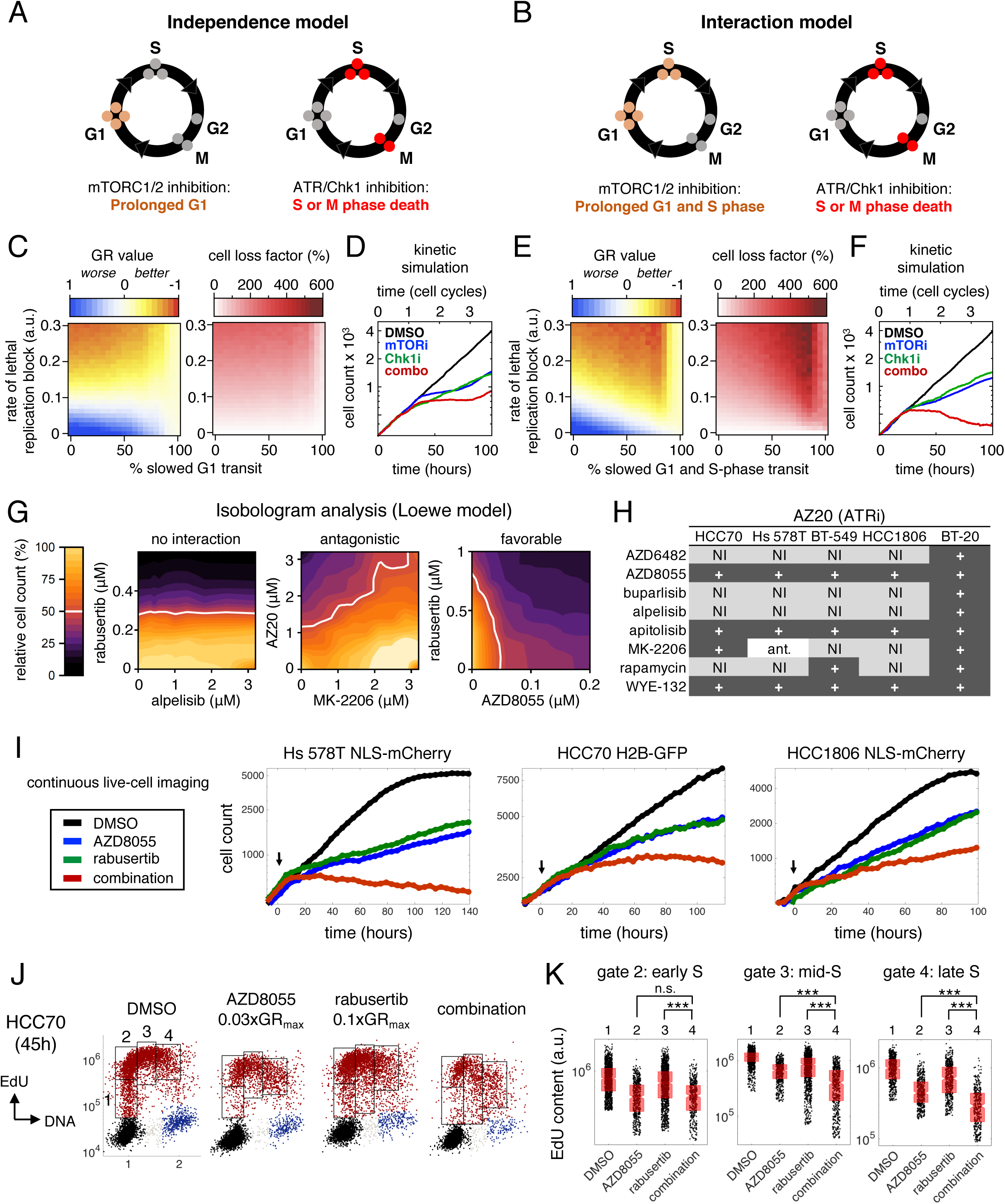
Combined inhibition of mTORC1/2 and PIKKs produces benefit by co-targeting pathways required in S phase. **A-B**. Schematics of the *independence* and *interaction* models. **C-D**. Simulated GR values (*left*), cell loss factor/cytotoxicity (*middle*) and growth curves (*right*) for the *independence* model, in which mTORC1/2 inhibition prolongs G1 and ATR/Chk1 inhibition causes lethality in S-phase or mitosis, as a consequence of blocked replication. **E-F**. Identical simulations as in C and D, but for the *interaction* model, in which mTORC1/2 inhibition prolongs both G1 and S-phase. **G.** Examples of no interaction, antagonism or favorable drug interaction by isobologram analysis and a Loewe dose equivalence model. White line indicates 50% reduction in viable cell count. **H**. Summary of interactions between PI3K pathway drugs and the ATR inhibitor AZ20. “+” indicates additive or greater benefit by Loewe criteria, “N.I.” indicates no interaction/independence, and “ant.” indicates antagonism. **I**. Live-cell imaging of cells treated with submaximal doses of AZD8055 and/or the Chk1 inhibitor rabusertib. Doses for each cell line (AZD8055, rabusertib): Hs 578T (100nM, 500nM); HCC70 (32nM, 320nM); HCC1806 (320nM, 320nM). **J**. DNA content vs EdU content at time=T_d_ (45h) for HCC70 cells treated with 0.03xGR_max_ dose (0.1μM) of AZD8055 and/or 0.1xGR_max_ dose (0.3μM) of rabusertib. Cells at S phase entry and in early, mid- and late S phase are gated (1-4). **K**. Total EdU content in S-phase cells in gates 2-4 in J. ***P<0.001 by Mann-Whitney U test; n.s.: not significant.

When the rate of cell killing caused by ATR/Chk1 inhibition was low (between 0 and 0.15 in **Figure 6C**), the *independence model* predicted greater cytostasis with increased mTORC1/2 inhibition. However, with higher cytotoxicity (values of 0.2 to 0.3), blocking mTORC1/2 had an antagonistic effect. A kinetic simulation of submaximal doses of an ATR/Chk1 inhibitor combined with an mTORC1/2 inhibitor demonstrated only a modest effect on viable cell number (**Figure 6D**). In contrast, the *interaction model* predicted substantially decreased GR values and increased cell killing across a wide range of activities of ATR/Chk1 and mTORC1/2 inhibitors (**Figure 6E**). Antagonism between mTORC1/2 and ATR/Chk1 inhibitors was observed only at the highest levels of mTORC1/2 inhibition, which markedly prolonged G1. Kinetic simulations of submaximal drug activities acting in combination yielded growth curves with negative slopes, denoting cytotoxicity (**Figure 6F**). Thus, the *interaction model* demonstrates that mTORC1/2 inhibitors causing incomplete G1/S block can be beneficially combined with ATR/Chk1 inhibitors if the combination increases the probability of S-phase cell killing. The cytotoxic effects of Torin2 and its analogs are thus likely to depend on the sensitivity of TNBC cells to both mTORC1/2 and PIKK inhibition during S phase.

To investigate whether combining PI3K/AKT/mTOR and ATR/Chk1 inhibitors might be broadly useful, we performed isobologram analysis on two-way dose-response landscapes for 16 different drug combinations—eight PI3K/AKT/mTOR drugs versus two ATR/Chk1 inhibitors—in five TNBC cell lines. Each of the 80 landscapes involved 100 different dose ratios. Using Loewe criteria (Tallarida, 2011), we classified combinations as exhibiting independence (no change in viable cell number compared to the more effective single agent) or favorable/antagonistic interactions (decreased/increased cell number compared to single agents) (**Figure 6G**). In total, 46 out of 80 combinations exhibited pharmacological interaction, of which 45 (98%) were at least additive and only one was antagonistic (**Figures 6H, S6A**). As predicted by modeling, favorable drug interactions gave rise to increased cell killing at submaximal concentrations of PI3K pathway inhibitors that only partly suppress mTORC1/2 signaling and incompletely block proliferation (**Figure 6I, S6B-C**). In HCC70 cells, for example, low doses of AZD8055 (0.1µM, 0.03xGR_max_) were sufficient to delay progression of S phase; when combined with low doses of rabusertib (0.32µM, 0.1xGR_max_), we observed significantly greater suppression of EdU incorporation in mid- and late S-phase cells compared to either drug alone (**Figures 6J-K**). Thus, as predicted by the *interaction model*, multiple PI3K pathway and ATR/Chk1 inhibitors can be beneficially combined in TNBC cell lines.

### Low doses of mTORC1/2 and ATR/Chk1 inhibitors in combination cause increased ssDNA in mitotic prophase and death

It remained unclear how low doses of mTORC1/2 and ATR/Chk1 inhibitors in combination cause increased cell killing. In HCC1806 cells, Torin2 exposure led to accumulation of large numbers of S_NR_ cells with reduced levels of geminin, cyclin A2 and p-pRb, and death by replication catastrophe (**Figures 2-4**). However, exposure of HCC70 cells to Torin2 resulted in substantially less accumulation of S_NR_ cells and no change in expression of geminin or cyclin A2. Thus, we hypothesized that the low dose combination might increase cytotoxicity in HCC70 cells by a mechanism distinct from replication catastrophe. ATR/Chk1 inhibitors used alone can induce mitotic entry of S-phase cells with damaged or incompletely-replicated DNA (King et al., 2015). To determine whether this phenomenon occurs in HCC70 cells treated with low-dose combinations, we examined early mitotic (prophase) cells in asynchronous drug-treated cultures, identified by positive staining for phospho-Histone H3 (p-HH3) (S10) and the presence of uncondensed chromatin (as visualized by Hoechst staining). To detect ssDNA, cells were labeled with BrdU for t=T_d_ and stained with anti-BrdU antibodies under native conditions (**Figure 7A**). We found that exposure to the combination of low-dose AZD8055 (0.1µM, 0.03xGR_max_) and rabusertib (0.32µM, 0.1xGR_max_) for t=0.5xT_d_ (24h) produced higher fractions of BrdU-positive prophase cells and higher nuclear BrdU intensity levels than either drug alone or than a 0.3xGR_max_ dose (1µM) of Torin2 (**Figures 7B-D**). Thus, the low-dose combination causes increased mitotic entry of HCC70 cells with ssDNA.

**Figure 7.**
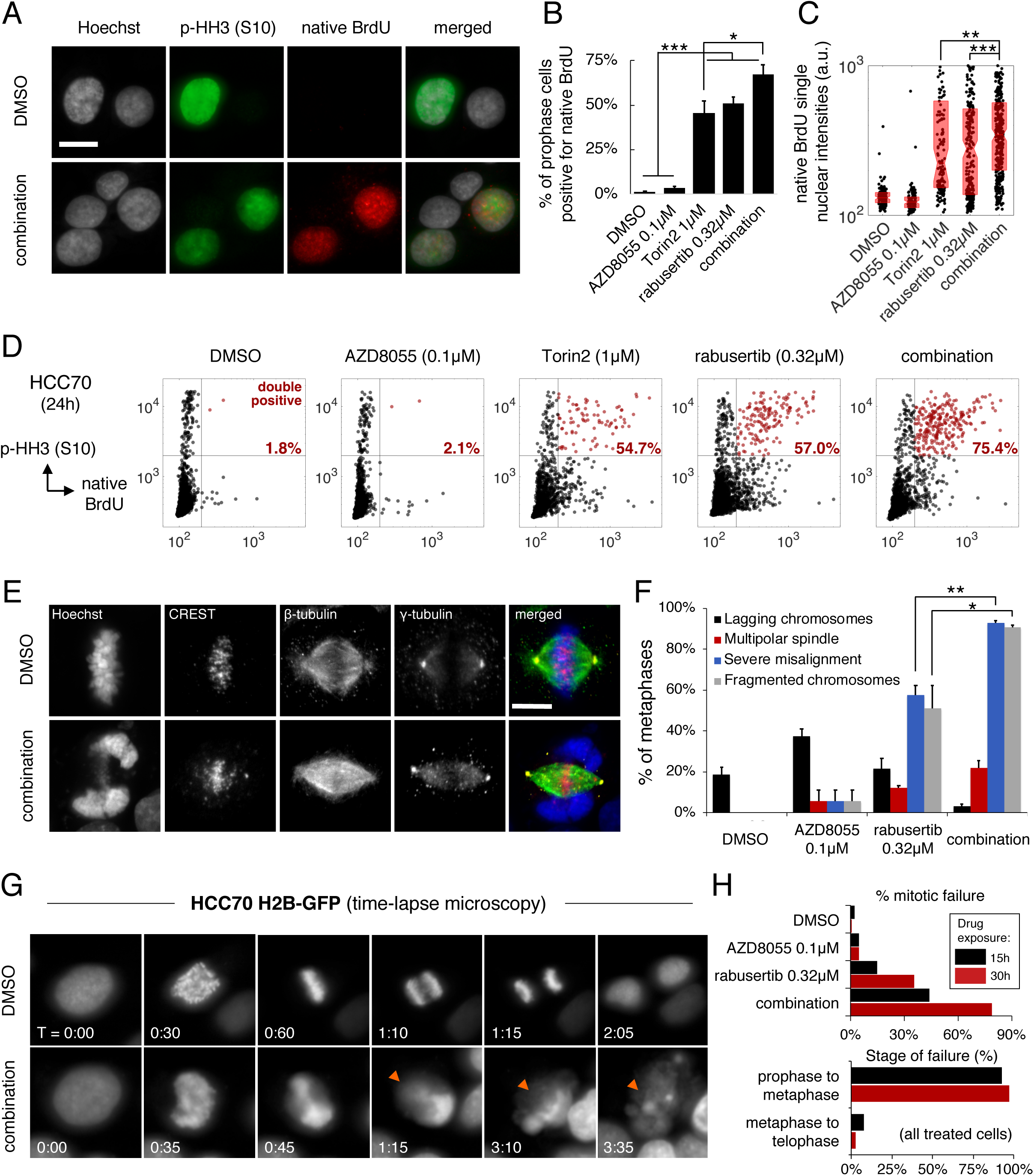
Low doses of mTORC1/2 and ATR/Chk1 inhibitors in combination cause increased ssDNA in mitotic prophase and death. “Combination” indicates 0.1μM AZD8055 and 0.32μM rabusertib throughout. **A.** Representative images of HCC70 cell nuclei after staining for Hoechst, p-HH3 and native BrdU after treatment for t=0.5xT_d_ (24h). Scale bar=20μM. **B**. Mean percent ± SEM of native BrdU/p-HH3 double-positive cells at 24h post-treatment; N=3 experiments. **C**. Native BrdU levels in single nuclei at 24h post-treatment. Boxplots show the median and 25th/75th percentiles. **D**. Representative scatterplots quantifying cells that are double-positive for p-HH3 and native BrdU (red) at 24h post-treatment, with percentages relative to the population of p-HH3-positive (prophase) cells. **E.** Maximum intensity projection of confocal image stacks of metaphase cells after exposure to DMSO control or the combination for 24h. Scale bar=10μm. **F**. Mean percent ± SEM of metaphases with the indicated aberrations at 24h post-treatment; N=2 experiments, 188 metaphases. **G**. Time-lapse imaging of HCC70 H2B-GFP cells showing aberrant chromatin condensation in cells treated with the combination, followed by abnormal metaphase (arrowhead) and death. Numbers on images denote hours:minutes. **H.** Percent of cells exhibiting mitotic failure at 15 or 30h post-treatment (*top*), and the timing of mitotic failure with respect to mitotic stage for all treated cells (*bottom*). N ∼200 cells at each timepoint. *P<0.05, **P<0.01, ***P<0.001, and n.s. (not significant) by ANOVA and Tukey’s test (panels **B, F**; only selected comparisons shown) and Mann-Whitney U-test (**C**).

To determine the fates of these cells, we stained chromosomes, centromeres, spindle fibers, and centrosomes using Hoechst and anti-CREST, anti-β-tubulin, and anti-γ-tubulin antibodies, following drug exposure for t=0.5xT_d_ (24h). Whereas AZD8055 alone decreased the total number of mitotic figures per high-powered field, rabusertib and the low-dose combination increased mitotic cell counts by 1.7-fold and 2.6-fold, respectively (N=672 fields; P<0.001 for combination vs rabusertib by one-way ANOVA and Tukey’s test) (**Figure S7A**). The combination also increased the percentage of mitotic cells in metaphase to 80%, compared to 28% for DMSO, 36% for AZD8055 alone and 58% for rabusertib alone (N=569 mitotic cells; P<0.01 for combination vs rabusertib) (**Figure S7B**). More than 90% of metaphase cells treated with AZD8055 plus rabusertib exhibited fragmented and/or misaligned chromosomes (N=188 metaphase cells; P<0.01 for combination vs rabusertib for severe misalignment and P<0.05 for chromosome fragmentation) (**Figures 7E-F**). Thus, cells with high levels of ssDNA are delayed in metaphase with incorrectly aligned or damaged chromosomes.

The interdependency of these events was made more obvious by time-lapse imaging of cells expressing H2B-GFP. Drug-induced abnormalities in chromatin condensation that were visible in prophase frequently led to aberrant metaphases (N=198 and 222 cells tracked through mitosis at 15 and 30h, respectively) (**Figure 7G**). Mitotic death ensued after varying lengths of time without evidence of progression to anaphase. Consistent with modeling data, the rate of mitotic entry was lower in cells treated with the low-dose combination versus rabusertib alone (0.8% vs 1.7% of cells per hour after 30h of drug exposure; **Figure S7C**). However, nearly 80% of cells entering mitosis in the presence of the drug combination underwent mitotic catastrophe (a 2.2-fold increase over rabusertib alone) (**Figure 7H**). Among all drug-treated cells that died during mitosis, >97% failed between prophase and metaphase. We conclude that low doses of mTORC1/2 inhibitors can augment the cytotoxicity of Chk1 inhibitors in TNBC by increasing the rate of mitotic failure.

## DISCUSSION

The high frequency of mutations in PI3K pathway genes across human cancers has motivated clinical development of numerous small-molecule kinase inhibitors (Janku et al., 2018), few of which have proven effective in patients. We find that the pre-clinical compound Torin2 is unusually active in blocking proliferation and inducing death of PI3K-activated, *TP53*-mutant TNBC cells. In contrast, Torin2 is only partially cytostatic and causes little or no death in non-transformed mammary epithelial cells. Torin2 was developed as an mTORC1/2 inhibitor, but we have found that its unusually high activity in TNBC results from combined inhibition of mTOR and structurally related PIKKs such as ATR. Torin2-like pharmacology can be reconstituted by mixing selective inhibitors of mTORC1/2 and PIKKs. Moreover, across a panel of Torin2 analogues, negative GR_max_ values indicative of cytotoxicity correlate with activity against both mTORC1/2 and PIKKs. In multiple TNBC lines, combinations of drugs targeting mTORC1/2 and ATR or Chk1, a checkpoint kinase that acts downstream of multiple PIKKs (Buisson et al., 2015), also show evidence of favorable drug interactions. Thus, the polypharmacology of Torin2 is essential for its high activity, and combined inhibition of mTORC1/2 and PIKKs may represent a novel strategy for targeting PI3K-activated TNBC.

The benefits of combining mTORC1/2 and ATR/Chk1 inhibitors are unexpected because these two drug classes primarily affect successive cell cycle states. PI3K/AKT/mTOR signaling canonically regulates the G1/S transition (Liang and Slingerland, 2003), while ATR/Chk1 is required during S phase and at the G2/M checkpoint (King et al., 2015; Zeman and Cimprich, 2014). Prior studies have shown that combinations of chemotherapies or chemotherapy plus radiation are antagonistic when successive cell cycle stages are affected (Johnson et al., 1999; Sui et al., 2004). This is intuitive, because cells blocked in G1 cannot proceed into later phases of the cell cycle where cytotoxicity occurs. However, in TNBC cells we find that PI3K pathway drugs also disrupt progression of S phase. Modeling and experimentation together suggest that combinations of mTORC1/2 and PIKK inhibitors can produce benefit under conditions in which mTORC1/2 inhibitors incompletely block cells at G1/S and yet slow the rate of replication in S phase. Thus, the relative insensitivity of TNBC cells to the anti-proliferative effects of mTORC1/2 inhibitors at G1/S is a liability for monotherapy, but an opportunity for combination therapies that exploit vulnerabilities in S phase. The strong effects of Torin2 on S-phase cells appear to result from a felicitous combination of reduced rates of DNA synthesis due to mTORC1/2 inhibition and reduced replication fork stability due to PIKK inhibition. The accumulation of ssDNA and DNA damage caused by exposure to Torin2 consequently result in death of TNBC cells in S phase or subsequent mitosis.

We find that the slowed rates of DNA synthesis caused by exposure of TNBC cells to PI3K pathway inhibitors are associated with altered levels of pyrimidine and purine nucleotides and their precursors. Prior studies have also identified roles for the PI3K pathway in regulating nucleotide biosynthesis: it promotes uptake and oxidative metabolism of glucose and directs metabolic flux through the non-oxidative arm of the pentose phosphate pathway. This increases levels of ribose 5-phosphate and phosphoribosylpyrophosphate (PRPP), which are required for synthesis of pyrimidines and purines (Juvekar et al., 2016b; Wang et al., 2009). The PI3K pathway also increases *de novo* synthesis of purines by regulating the bifunctional enzyme 5-aminoimidazole-4-carboxamide ribonucleotide formyltransferase/IMP cyclohydrolase (Wang et al., 2009). mTORC1 is known to regulate SREPB1 and transcription of enzymes involved in glycolysis and the oxidative arm of the pentose phosphate pathway (Düvel et al., 2010); CAD activity, which catalyzes *de novo* pyrimidine synthesis; and expression of *MTHFD2*, which increases *de novo* purine biosynthesis (Ben-Sahra et al., 2013, 2016). In TNBC cells, we find that acute inhibition of mTORC1 also causes increased levels of cytidine and cytosine, known substrates for pyrimidine salvage pathways, as well as changes in the levels of dTDP and dTMP consistent with reduced activity of deoxythymidylate kinase, an enzyme that acts downstream of both *de novo* synthesis and salvage pathways for pyrimidines. Exposure of cells to mTOR inhibitors may also perturb S phase by additional mechanisms. For example, mTORC1/2 regulates the transcription and translation of multiple genes and proteins required for DNA replication and repair in response to DNA damage and replication stress (Silvera et al., 2017).

Dependencies on ATR/Chk1 signaling for S-phase progression are also likely to be multifactorial in origin (Zeman and Cimprich, 2014). Oncogenes commonly amplified in TNBC, such as *MYC* and *CCNE1*, cause excessive origin firing and conflicts between replication and transcription, resulting in stalled replication forks (Macheret and Halazonetis, 2018). Stalled forks generate ssDNA due to uncoupling of replicative helicases and DNA polymerases, which activates ATR/Chk1 signaling (Zeman and Cimprich, 2014). Inhibition of ATR/Chk1 in this setting causes accumulation of ssDNA, consumption of replication factors, DNA damage and “replication catastrophe” (Toledo et al., 2013). Loss-of-function mutations in *TP53* may also be important, since absence of the DNA damage checkpoint at G1/S renders cells more dependent on ATR/Chk1 signaling for the function of alternative cell cycle checkpoints to maintain genomic integrity (Ma et al., 2011). By inhibiting both mTORC1/2 and PIKKs, Torin2 is therefore able to effectively exploit S-phase and checkpoint vulnerabilities in TNBC to promote tumor cell killing.

### Translational prospects

The foremost challenge in targeting the PI3K pathway for anti-cancer therapy is achieving anti-proliferative effects and cell killing sufficient to block tumor growth at doses tolerated by patients. The dual PI3K/mTOR inhibitor omipalisib is emblematic of this challenge: omipalisib produces substantial growth inhibition and apoptosis in TNBC cell lines, but the maximum tolerated dose in patients fails to fully inhibit intratumoral AKT and has only modest anti-tumor activity (Munster et al., 2016). To improve the therapeutic index, recent efforts have focused on development of selective inhibitors of PI3K p110 isoforms (α, β, δ, γ) but these drugs often lack sufficient activity in solid tumors as monotherapy (Costa et al., 2015; Elkabets et al., 2013; Schwartz et al., 2015). In this context, the polypharmacology of Torin2-like compounds appears promising, in that it eliminates the requirement for strong inhibition of mitogenic signaling and G1/S arrest to produce efficacy. We have also shown that PI3K pathway inhibitors could be used at relatively low doses in combination with other drugs to exploit replicative and/or checkpoint vulnerabilities and produce cytotoxicity. Thus, new combinatorial strategies inspired by Torin2 may not only be more effective, but also have a greater therapeutic index in patients.

## AUTHOR CONTRIBUTIONS

SSC led the study; SSC, AJ, MC and CM performed experiments; SSC analyzed the data; AP produced the models and isobolograms; JMA, MN, CY and JC contributed to experimental design; QL and SCS assisted with chemistry and NG and PKS supervised the research. All authors have reviewed and approved the entire content of this submission.

## ACKNOWLEDGEMENTS

We thank U Matulonis for supporting this work and M Hafner, M Yuan and A Chen for their assistance with experiments and the manuscript.

## STAR METHODS

Further information and requests for resources and reagents should be directed to and will be fulfilled by the Lead Contact, Peter Sorger.

## EXPERIMENTAL MODEL AND SUBJECT DETAILS

### Cell lines

The media and culture conditions for all breast cancer and non-transformed mammary epithelial cell lines used in this study are described below. All cell lines were isolated from females, to the best of our knowledge. STR profiling was performed to confirm cell line identity.

| Cell line | RRID | Growth medium<br>(FBS: heat-inactivated fetal bovine serum;<br>P/S: penicillin/streptomycin) | Temp | CO<sub>2</sub> |
| --- | --- | --- | --- | --- |
| **184-B5** | CVCL_4688 | MEBM (Lonza/Clonetics; MEGM Kit #CC-3150) + 1ng/ml cholera toxin (unfiltered) | 37°C | 5% |
| **AU-565** | CVCL_1074 | RPMI-1640 + 10% FBS + 1% P/S | 37°C | 5% |
| **BT-20** | CVCL_0178 | EMEM + 10% FBS + 1% P/S | 37°C | 5% |
| **BT-474** | CVCL_0179 | RPMI-1640 + 10% FBS + 1% P/S | 37°C | 5% |
| **BT-483** | CVCL_2319 | RPMI-1640 + 20% FBS + 1% P/S + 0.01mg/ml bovine insulin | 37°C | 5% |
| **BT-549** | CVCL_1092 | RPMI-1640 + 10% FBS + 1% P/S + 0.023IU/ml insulin | 37°C | 5% |
| **CAL-120** | CVCL_1104 | DMEM + 10% FBS + 1% P/S | 37°C | 5% |
| **CAL-51** | CVCL_1110 | DMEM + 20% FBS + 1% P/S | 37°C | 5% |
| **CAL-85-1** | CVCL_1114 | DMEM + 10% FBS + 1% P/S | 37°C | 5% |
| **CAMA-1** | CVCL_1115 | EMEM + 10% FBS + 1% P/S | 37°C | 5% |
| **HCC1143** | CVCL_1245 | RPMI-1640 + 10% FBS + 1% P/S | 37°C | 5% |
| **HCC1395** | CVCL_1249 | RPMI-1640 + 10% FBS + 1% P/S | 37°C | 5% |
| **HCC1419** | CVCL_1251 | RPMI-1640 + 10% FBS + 1% P/S | 37°C | 5% |
| **HCC1428** | CVCL_1252 | RPMI-1640 + 10% FBS + 1% P/S | 37°C | 5% |
| **HCC1500** | CVCL_1254 | RPMI-1640 + 10% FBS + 1% P/S | 37°C | 5% |
| **HCC1569** | CVCL_1255 | RPMI-1640 + 10% FBS + 1% P/S | 37°C | 5% |
| **HCC1806** | CVCL_1258 | RPMI-1640 + 10% FBS + 1% P/S | 37°C | 5% |
| **HCC1937** | CVCL_0290 | RPMI-1640 + 10% FBS + 1% P/S | 37°C | 5% |
| **HCC1954** | CVCL_1259 | RPMI-1640 + 10% FBS + 1% P/S | 37°C | 5% |
| **HCC202** | CVCL_2062 | RPMI-1640 + 10% FBS + 1% P/S | 37°C | 5% |
| **HCC38** | CVCL_1267 | RPMI-1640 + 10% FBS + 1% P/S | 37°C | 5% |
| **HCC70** | CVCL_1270 | RPMI-1640 + 10% FBS + 1% P/S | 37°C | 5% |
| **Hs 578T** | CVCL_0332 | DMEM + 10% FBS + 1% P/S | 37°C | 5% |
| **hTERT-HME1** | CVCL_3383 | MEBM + Lonza CC-3150 kit | 37°C | 5% |
| **MCF-10A** | CVCL_0598 | 1:1 DMEM and Ham's F12 + 20ng/ml human epidermal growth factor + 100ng/ml cholera toxin + 0.01mg/ml bovine insulin + 500ng/ml hydrocortisone, 95% + 5% horse serum | 37°C | 5% |
| **MCF-10F** | CVCL_3633 | 1:1 DMEM and Ham's F12 + 20ng/ml human epidermal growth factor + 100ng/ml cholera toxin + 0.01mg/ml bovine insulin + 500ng/ml hydrocortisone, 95% + 5% horse serum | 37°C | 5% |
| **MCF-12A** | CVCL_3744 | 1:1 DMEM and Ham's F12 + 20ng/ml human epidermal growth factor + 100ng/ml cholera toxin + 0.01mg/ml bovine insulin + 500ng/ml hydrocortisone, 95% + 5% horse serum | 37°C | 5% |
| **MCF-7** | CVCL_0031 | DMEM + 10% FBS + 1% P/S | 37°C | 5% |
| **MDA-MB-134-VI** | CVCL_0617 | Leibovitz's L-15 + 20% FBS (not heat-inactivated) + 1% P/S | 37°C | 0% |
| **MDA-MB-157** | CVCL_0618 | Leibovitz's L-15 + 10% FBS (not heat-inactivated) + 1% P/S | 37°C | 0% |
| **MDA-MB-175-VII** | CVCL_1400 | Leibovitz's L-15 + 10% FBS (not heat-inactivated) + 1% P/S | 37°C | 0% |
| **MDA-MB-231** | CVCL_0062 | DMEM (or L-15) + 10% FBS (not heat-inactivated) + 1% P/S | 37°C | 0% |
| **MDA-MB-361** | CVCL_0620 | Leibovitz's L-15 + 20% FBS (not heat-inactivated) + 1% P/S | 37°C | 0% |
| **MDA-MB-415** | CVCL_0621 | Leibovitz's L-15 + 15% FBS (not heat-inactivated) + 1% P/S + 2mM L-glutamine + 10mg/l insulin | 37°C | 0% |
| **MDA-MB-436** | CVCL_0623 | Leibovitz's L-15 + 10% FBS (not heat-inactivated) + 1% P/S + 10mg/l insulin | 37°C | 0% |
| **MDA-MB-453** | CVCL_0418 | Leibovitz's L-15 + 10% FBS (not heat-inactivated) + 1% P/S | 37°C | 0% |
| **MDA-MB-468** | CVCL_0419 | Leibovitz's L-15 + 10% FBS (not heat-inactivated) + 1% P/S | 37°C | 0% |
| **SK-BR-3** | CVCL_0033 | McCoy's 5a + 10% FBS + 1% P/S | 37°C | 0% |
| **SUM102PT** | CVCL_3421 | Ham's F-12 + 1% P/S + 5 µg/ml insulin + 1ug/ml hydrocortisone + 10ng/mL epidermal growth factor + 5mM ethanolamine + 10mM HEPES + 5ug/mL transferrin + 10nM T3 + 50nM sodium selenite + 1g/L BSA | 37°C | 5% |
| **SUM1315MO2** | CVCL_5589 | Ham's F-12 + 5% FBS + 1% P/S + 10ng/ml human epidermal growth factor + 5µg/ml insulin + 10 mM HEPES | 37°C | 5% |
| **SUM149PT** | CVCL_3422 | Ham's F-12 + 5% FBS + 1% P/S + 1ug/ml hydrocortisone + 5µg/ml insulin + 10mM HEPES | 37°C | 5% |
| **SUM159PT** | CVCL_5423 | Ham's F-12 + 5% FBS + 1% P/S + 1ng/ml hydrocortisone + 5µg/ml insulin + 10mM HEPES | 37°C | 5% |
| **SUM225CWN** | CVCL_5593 | Ham's F-12 + 5% FBS + 1% P/S + 5µg/ml insulin + 1ug/ml hydrocortisone + 10mM HEPES | 37°C | 5% |
| **SUM44PE** | CVCL_3424 | Ham's F-12 + 1% P/S + 5µg/ml insulin + 1ug/ml hydrocortisone + 5mM ethanolamine + 10mM HEPES + 5ug/mL transferrin + 10nM T3 + 50nM sodium selenite + 1g/L BSA | 37°C | 5% |
| **SUM52PE** | CVCL_3425 | Ham's F-12 + 5% FBS + 1% P/S + 5 µg/ml insulin + 1ug/ml hydrocortisone + 10mM HEPES | 37°C | 5% |
| **T-47D** | CVCL_0553 | RPMI-1640 + 10% FBS + 1% P/S + 0.2units/ml bovine insulin | 37°C | 5% |
| **UACC-812** | CVCL_1781 | Leibovitz's L-15 + 20% FBS (not heat-inactivated) + 1% P/S + 2mM L-glutamine + 20ng/ml human EGF | 37°C | 0% |
| **UACC-893** | CVCL_1782 | Leibovitz's L-15 + 10% FBS (not heat-inactivated) + 1% P/S | 37°C | 0% |
| **ZR-75-1** | CVCL_0588 | RPMI-1640 + 10% FBS + 1% P/S | 37°C | 5% |
| **ZR-75-30** | CVCL_1661 | RPMI-1640 + 10% FBS + 1% P/S | 37°C | 5% |

### METHOD DETAILS

### Western blotting

To determine protein levels under unstimulated conditions, cells grown to approximately 75% confluency under the conditions described above were serum starved overnight and then lysed with Mammalian Protein Extraction Buffer (M-PER; Thermo Scientific, 78501) supplemented with protease inhibitor cocktail (Sigma-Aldrich, P2714), 1mM sodium orthovanadate (Sigma-Aldrich, S6508), 5mM sodium pyrophosphate (Sigma-Aldrich, 221368), 50μM oxophenylarsine (EMD Biosciences, 521000) and 10mM bpV (phen) (EMD Biosciences, 203695). Lysed cells were scraped off the plate, collected in microcentrifuge tubes and incubated on ice for 30min. Membranes and cell debris were sedimented by centrifugation at 20,000x*g* for 10min at 4°C. Supernatants were pooled, aliquoted and stored at −80°C. Protein concentration was determined using the Pierce BCA protein assay kit (Thermo Fisher Scientific, 23225). Protein electrophoresis was performed using E-PAGE pre-cast 8% gels with 48 wells (Thermo Fisher Scientific, EP04808). Separated protein was transferred to Immobilon-FL PVDF membranes (EMD Millipore, IPFL00005) using the Criterion blotting system (Bio-Rad). Membranes were blocked for 1h at room temperature (RT) with Odyssey blocking buffer (OBB; Li-COR 927-40000), incubated in primary antibodies in OBB overnight at 4°C, washed for 30min in PBS plus 0.1% Tween-20 (PBST), incubated in secondary antibody in OBB for 1h at RT, and washed again for 30min in PBST. Anti-PTEN (Cell Signaling Technologies, 9188) and anti-INPP4B (Epitomics, 2512-1) primary antibodies were each used at 1:1000 dilution. Detection was performed using anti-rabbit IRDye 800CW pre-adsorbed secondary antibody (Rockland, 613-431-028) at 1:20,000 dilution. The fluorescence intensity of bands was quantified using the Odyssey system (Li-COR).

### Evaluation of drug sensitivity

All compounds except Torin2 analogs were obtained from MedChem Express. Stock solutions were prepared at a concentration of 10mM in DMSO and stored at -30°C. Torin2 analogs (QL-VIII-58, QL-V-107, QL-IV-100, QL-VI-86, QL-XII-47, QL-V-73, QL-X-138, QL-XII-108, QL-XII-61) were obtained from the laboratory of Nathanael Gray (Dana-Farber Cancer Institute) as 10mM stock solutions in DMSO. PI3K pathway drugs were arrayed in master library plates and pin-transferred to cells in Cell Carrier 384 well plates (Perkin Elmer, 6007550). Torin2 analogs were delivered to cells in Cell Carrier 384 well plates using the D300 digital dispenser and accompanying software (Hewlett Packard). Cells were plated at a density of 1,000-2,000 cells per well using the Multidrop Combi (Thermo Scientific) and treatments were performed on the following day approximately 18-24h after plating. Inhibitors were administered at 9 different concentrations in at least technical triplicate and DMSO alone was used as a control. To determine cell number at the start (time = 0) and end (t = 72h) of treatment, cells were arrayed in two different plates with the time = 0 plate receiving no treatment and the 72h plate receiving DMSO control and active drug. Viable cells were identified using 15μL of staining solution, comprising 1:5,000 LIVE/DEAD Far Red Dead Cell Stain (Invitrogen, L10120), 1:10,000 Hoechst 33342 trihydrochloride trihydrate 10mg/mL solution in water (Invitrogen, H3570) and 1:10 OptiPrep (Sigma-Aldrich, D1556) in PBS. Cells were incubated in staining solution for 30min at 37°C with 5% CO_2_ and then fixed with 20μL fixing solution, comprising 1:20 37% formaldehyde (Sigma-Aldrich, F1635) and 1:5 OptiPrep in PBS, for 30min at RT. To achieve accurate cell counts, no aspiration of well contents was performed prior to fixing cells. Following fixation, all reagents were aspirated and replaced with 60μL of PBS using an EL406 Washer Dispenser (BioTek). Cells were then imaged using the Operetta High-Content Imaging System (Perkin Elmer). Six fields of view spanning each well in its entirety were acquired using the 10x high-NA objective lens and appropriate excitation/emission filters for Hoechst and LIVE/DEAD Far Red cell stains. The total viable cell count per well was determined using Columbus image analysis software (Perkin Elmer), as described below.

### Immunofluorescence microscopy

Cells were plated in Cell Carrier 384 well plates at either 1,000 (Hs 578T) or 2,000 (BT-20, BT-549, HCC1806, HCC38, HCC70) cells per well, treated with active drug or DMSO using the D300 digital dispenser, and fixed at various timepoints using 4% formaldehyde in PBS for 30min at RT. Cells were washed twice with PBS for 5min each at RT using the EL406 automated plate washer, permeabilized with 0.25% Triton X-100 (Bio-Rad, 1610407) in PBS for 15min, washed twice again for 5min each with PBS and blocked with OBB for at least 1h at RT. Cells were incubated with primary antibody diluted in OBB overnight at 4**°**C. The following morning, cells were washed three times with PBST for 5min each and then stained with secondary antibodies diluted 1:2,000 in OBB for 1h at RT. Secondary antibodies varied for different experiments but included Alexa Fluor 647 donkey anti-rabbit (Invitrogen, A31573), Alexa Fluor 647 donkey anti-mouse (Invitrogen, A31571), Alexa Fluor 647 goat anti-human (Invitrogen, A21445), Alexa Fluor 488 donkey anti-rabbit (Invitrogen, A21206), Alexa Fluor 488 goat anti-mouse (Invitrogen, A11001), and Alexa Fluor 568 goat anti-rabbit (Invitrogen, A11011). Cells were washed twice with PBST and once with PBS for 5min each prior to staining with Hoechst 33342 (Thermo Scientific, 1:10,000 in PBS) for 30min at RT. To facilitate segmentation of signal intensities specifically in the nucleus, cytoplasm, or whole cell in certain experiments, cells were simultaneously stained with 1:5000 Cellomics Whole Cell Stain (Thermo Scientific, 8403502) in PBS. Cells were washed twice with PBS, sealed with foil, and imaged using the Operetta High-Content Imaging System, the ImageXpress Micro Confocal High-Content Imaging System (Molecular Devices) or the IN Cell Analyzer 6000 (GE Healthcare Life Sciences) using the appropriate excitation and emission filters, depending on the specific assay.

### Primary antibodies and dilutions

The following primary antibodies were obtained from Cell Signaling Technology (catalog number, dilution in OBB): rabbit anti-phospho-AKT T308 (13038, 1:400), rabbit anti-phospho-AKT S473 (4060, 1:200), rabbit anti-phospho-Chk1 S317 (12302, 1:800), rabbit anti-phospho-Chk2 T68 (2661, 1:100), rabbit anti-phospho-4E-BP1 T37/46 (2855, 1:200), rabbit anti-phospho-S6 S235/236 (4858, 1:1000), rabbit anti-phospho-H2A.X S139 (9718, 1:400), rabbit anti-phospho-Histone H3 S10 (3377, 1:800), mouse anti-β-tubulin (86298, 1:400), rabbit anti-β-tubulin (2128, 1:400), mouse anti-BrdU (Bu20a) (5292, 1:200), rabbit anti-phospho-GSK3B-S9 (5558, 1:400), rabbit anti-FOXO3 (2497, 1:200), rabbit anti-p21 (2947, 1:400), rabbit anti-p27 (3686, 1:400), and rabbit anti-phospho-Rb S807/811 (8516, 1:800). The following primary antibodies were obtained from Abcam (catalog number, dilution in OBB): rabbit anti-cyclin D1 (ab134175, 1:100), rabbit anti-geminin (ab195047, 1:100), and rabbit anti-cyclin A2 (ab181591, 1:500). Rabbit anti-phospho-RPA-S4/8 antibodies were obtained from Bethyl (A300-245A, 1:400), rabbit anti-γ-tubulin antibodies from Sigma (T5192, 1:400), and Human Antibody against Centromere (CREST) antibodies from ImmunoVision (HCT-0100, 1:1000).

### Single-stranded DNA (ssDNA) and cell cycle analysis

To measure ssDNA, newly-synthesized DNA in proliferating cells was first labeled with 10µM BrdU (Sigma, B9285) in growth media for a time equal to the measured cell line division time (T_d_) in culture. Drug treatments were administered to labeled cells at T_d_ using the D300 digital dispenser, and cells were fixed at various timepoints using 4% formaldehyde. BrdU was detected under non-denaturing (“native”) conditions by standard immunofluorescence microscopy, as described above. Cell cycle analysis was performed after treating cells with active drug or DMSO-only control for time=T_d_, which was 28h for HCC1806 cells, 32h for Hs 578T cells, 40h for BT-549 cells, 45h for both HCC70 and HCC38 cells and 48h for BT-20 cells. Active S-phase cells were labeled with 10µM EdU (Thermo Fisher Scientific, C10637) in growth media for 0.025xT_d_ in an incubator. Cells were then fixed with 4% formaldehyde in PBS for 30min at RT. Incorporated EdU was detected using the Click-iT EdU Plus Alexa Fluor 488 imaging kit (Thermo Fisher Scientific, C10637), per the manufacturer’s instructions. Following the Click-iT reaction, cells were counterstained with 5μg/mL Hoechst dye and imaged using the Operetta or ImageXpress Micro Confocal High-Content Imaging System. In certain experiments, EdU-labeled cells were counterstained with anti-geminin, anti-cyclin A2, anti-phospho-pRb or anti-phospho-H2A.X antibodies overnight following the Click-iT reaction and prior to Hoechst staining. Continuous labeling of S-phase cells was performed using 1µM F-ara-EdU (Sigma, T511293) for 48 or 75h.

### Live cell microscopy

Time-lapse imaging of proliferation and apoptosis in live cells was performed using the Incucyte Zoom live-cell analysis system (Essen Bioscience) inside of an incubator maintained at 37**°**C and 5% CO_2_. Cells were plated at low density in 384-well plates (Corning, 3712) and images were taken of whole wells every 30-120min using the 4x lens. Drugs were administered during logarithmic-phase growth. For proliferation assays, stable cell lines with nuclear-expressing fluorophores were used to quantify live cell number at regular time intervals. HCC70 H2B-GFP cells were established via lentiviral transduction (Addgene, 25999) followed by FACS sorting. The generation of Hs 578T NLS-mCherry and HCC1806 NLS-mCherry cell lines, which also express a F3aN400-Venus construct not utilized here, were previously described (Sampattavanich et al., 2018). Apoptotic cells were identified in wild-type cell lines using the Incucyte caspase 3/7 (C3/7) apoptosis assay reagent (Essen Bioscience, 4440), which was added to growth media at the recommended final concentration (5µM) several hours in advance of drug treatment. For each experimental replicate, the number of apoptotic cells was measured for each drug, dose, timepoint and cell line in technical triplicate.

For live-cell studies of mitosis, HCC70 H2B-GFP cells were plated at 2,000 cells per well in Cell Carrier 384-well plates. Plates were kept in a standard tissue culture incubator maintained at 37**°**C and 5% CO_2_ for approximately 48h before cells were treated with drugs using the D300 digital dispenser. Starting at either 15 or 30h after drug treatment, cells were imaged every 5-10 minutes for 5-7h in a humidified chamber maintained at 37**°**C and 5% CO_2_ using the 40x/0.95 NA lens and the FITC filter set of the IN Cell Analyzer 6000 microscope. The rate of mitotic entry and the percentage and stage of aberrations were visually scored for individual cells followed from prophase through cytokinesis in time-lapse image stacks.

To analyze the effects of drugs on cells in different stages of the cell cycle, HCC1806 cells expressing H2B-mTurquoise, mVenus-hGeminin(1-110) and mCherry-dE2F1N were established via sequential lentiviral transduction of three different plasmids (pCSII-EF-H2B-mTurquoise, pCSII-EF-Venus-geminin(1-110), and pCSII-EF-mCherry-dE2F1N), followed by fluorescence-activated cell sorting (FACS) for mTurquoise/mVenus double-positive cells, expansion, and a second FACS sort for mCherry-positive cells. HCC1806 triple-reporter cells were plated at 4,000 cells per well in 96-well plates (Ibidi, 89626), kept in a standard tissue culture incubator overnight, treated with drugs the next morning using the D300 digital dispenser and then imaged every 12min for 48h in a humidified chamber maintained at 37**°**C and 5% CO_2_ using the 20x/0.75NA lens of the ImageXpress Micro high-content imaging system with binning 2x2. mTurquoise, mVenus, and mCherry were excited using a Spectra Gen III (Lumencor) with 438/29nm, 510/25nm, and 578/21nm filters. Emission was collected using 482/25nm, 549/12nm, 623/32nm filters. To minimize the effect of phototoxicity, cells were imaged at minimal laser intensity and exposure times necessary to achieve approximately 8-fold actual signal intensity versus background for each fluorophore. At the end of the experiment, cells were stained with Hoechst and LIVE/DEAD stain to identify and quantify viable cells, as described above. To assess for possible phototoxicity, cell counts for DMSO-only control wells on the same plate that did not undergo imaging were compared to cell counts for DMSO-only control wells that underwent time-lapse imaging. To validate environmental conditions such as temperature, humidity, and CO_2_, viable cells in an identically-treated plate kept in a standard tissue culture incubator set at 37**°**C and 5% CO_2_ for 48h (*i.e*., the duration of the experiment) were also compared by Hoechst and LIVE/DEAD co-staining.

### Extraction of polar metabolites

Measurement of intracellular polar metabolite levels was performed to detect drug-induced changes in metabolic activity including *de novo* nucleotide biosynthesis and salvage pathways. Because nucleotides are synthesized primarily in S phase, drug-induced cell cycle changes have the potential to strongly bias metabolite levels. BT-549 cells were selected for profiling after acute drug exposure (*i.e.*, 6h) because we observed clear inhibition of DNA synthesis in S-phase cells without any detectable changes in the overall distribution of cells in different cell cycle stages. Cells in complete media were plated in 10cm^2^ dishes at 2.5x10^6^ cells per dish and cultured at 37°C and 5% CO_2_. Approximately two days later when cells were 75-80% confluent, the media was replaced with 10mL of fresh media containing rapamycin, AZD8055 or Torin2, or an equivalent volume of DMSO. After 6h of treatment, the 10cm^2^ dishes were placed on dry ice and the media was aspirated and replaced with ice cold 80% (v/v) methanol. Dishes were kept at -80°C for 20min prior to using a cell scraper to produce a cell lysate/methanol mixture. The mixture for each sample was collected in a 15mL Eppendorf tube and cellular debris and protein were pelleted by centrifugation at 1,800xg for 5min at 4°C. The methanol supernatant containing polar metabolites was transferred to a new 15mL tube on dry ice and the cell debris/protein pellets were resuspended two additional times with methanol, centrifuged and re-pelleted. Recovered methanol supernatants for each sample were pooled and then divided equally into four 1.5mL tubes on dry ice. Methanol was evaporated using a SpeedVac (no applied heat) and the resulting polar metabolite pellets were stored at -80°C.

### Metabolomics profiling by targeted mass spectrometry

Metabolites were re-suspended in 20mL HPLC-grade water for mass spectrometry. 5-7μL were injected and analyzed using a hybrid 5500 QTRAP triple quadrupole mass spectrometer (AB/SCIEX) coupled to a Prominence UFPLC system (Shimadzu) via selected reaction monitoring (SRM) of a total of 262 endogenous water-soluble metabolites for steady-state analyses of samples. Some metabolites were targeted in both positive and negative ion mode for a total of 298 SRM transitions using positive/negative ion polarity switching. ESI voltage was +4950V in positive ion mode and –4500V in negative ion mode. The dwell time was 3msec per SRM transition, and the total cycle time was 1.55sec. Approximately 10-14 data points were acquired per detected metabolite. Samples were delivered to the mass spectrometer via hydrophilic interaction chromatography (HILIC) using a 4.6mm i.d x 10cm Amide XBridge column (Waters) at 400 μL/min. Buffer A was comprised of 20mM ammonium hydroxide/20mM ammonium acetate (pH=9.0) in 95:5 water:acetonitrile and Buffer B was HPLC grade acetonitrile. Gradients were run starting from 85% buffer B (in buffer A) to 42% B from 0-5min; 42% B to 0% B from 5-16min; 0% B was held from 16-24min; 0% B to 85% B from 24-25min; 85% B was held for 7min to re-equilibrate the column. Peak areas from the total ion current for each metabolite SRM transition were integrated using MultiQuant v2.1 software (AB/SCIEX). Data were analyzed using MetaboAnalyst 4.0, as described below.

### Isobolograms and analysis of drug interactions

Viable cell counts were determined (as described above) after exposure to PI3K pathway and ATR/Chk1 inhibitors administered in combinations over two-dimensional gradients. Dose ranges typically spanned at least one order of magnitude and were selected based on the GR50 values for each single agent. Continuous interpolations were constructed over the 2D response surface, which were then plotted as contour maps that highlight contours of equal inhibitory effect, or isoboles (Interpolation function and ContourPlot functions in Mathematica v11). As described by the Loewe additivity model, drug combinations were classified as “additive” based on the concept of dose equivalence, which manifests as diagonal isoboles that intersect both axes. Combinations were classified as “antagonistic” when isoboles were observed to deviate away from either axis, indicating that the drugs used together were less effective than either drug used alone. Combinations were classified as having “no interaction” when the activity of each drug appeared to be unaffected by the second drug, indicating independent action and manifested by linear isoboles. Examples of each case are shown in **Figure 6G**.

### Cell cycle model and simulation of combinatorial drug effect

The effects of mTORC1/2 or ATR/Chk1 inhibition (individually or in combination) were simulated by a discrete time-step Monte Carlo simulation. This simulation followed the progress of individual cells through the G1, S, G2, and M phases of the cell cycle, with each cell possessing a cell-state variable that increments with cell cycle progression. When a cell completed M phase, the mother cell was replaced by two daughter cells at the start of G1. Simulations were initialized with a population of 300 cells distributed throughout the cell cycle (*i.e.*, asynchronous) based on experimentally measured cell cycle distributions for TNBC. mTORC1/2 inhibition slowed each cell’s progress through G1 (*independence* model) or through both G1 and S phases (*interaction* model) by imposing a lower probability per time-step that the cell-state variable will increase. ATR/Chk1 inhibition imposed on each S-phase cell a specific rate of lethal replication block at each time-step. In light of prior knowledge that the effects of ATR/Chk1 inhibition during replication can ultimately cause death of cells in S phase (replication catastrophe) or in mitosis (mitotic catastrophe), simulations did not discriminate between whether the moment of death arises in S phase or in mitosis. In the *interaction* model, slowing of cells through S phase by mTORC1/2 inhibition was found to elicit an increase in the accumulated probability of lethal damage caused by ATR/Chk1 inhibition.

### QUANTIFICATION AND STATISTICAL ANALYSIS

Details regarding the number of experimental replicates, specific statistical tests used and the level of significance are described in the primary and supplementary figure legends. Significant differences are also indicated on the primary and supplemental figures.

### Viable cell counts

Images of TNBC cells co-stained with Hoechst and LIVE/DEAD stains were analyzed using Columbus image data storage and analysis system (Perkin Elmer) in order to quantify the total number of viable cells per well. Nuclear segmentation was performed using the Hoechst fluorescence intensity. For example, the number of HCC1806 cell nuclei in a particular well was determined using the following segmentation routine: Module: ’Find Nuclei’; Channel: ‘Hoechst’; Method: M; diameter 28μm; splitting coefficient: 0.45; common threshold: 0.05. Because of differences in nuclear size and morphology, optimal segmentation routines varied for each TNBC cell line and were determined in an empirical fashion. After identification of all nuclei in a well, additional features were extracted from images to exclude over/under segmented nuclei, doublets and non-viable cells from the total cell counts. Over/under segmented nuclei and doublets were of atypical size and shape compared to appropriately segmented nuclei. Non-viable cells were characterized by positive LIVE/DEAD staining and/or small pyknotic nuclei with extremely bright Hoechst signal. Collectively, the image analysis features that best identified these characteristics included nuclear area and roundness (module: ’Calculate Morphology Properties’); nuclear Hoechst intensity (module: ’Calculate Intensity Properties’); nuclear LIVE/DEAD stain intensity (module: ’Calculate Intensity Properties’); Hoechst texture features (module: ’Calculate Texture Properties’; SER Features, spot; scale: 4px; normalization by: unnormalized); and LIVE/DEAD texture features (module: ’Calculate Texture Properties’; SER Features; scale: 6px; normalization by: kernel). These extracted features were then used to set filters (*i.e*., gates) in the module ’Select Population.’ In order to exclude extreme outliers representing non-viable or incorrectly segmented cells, filter settings were tuned interactively for each experimental replicate by visually comparing images of cells in DMSO-treated wells against images of cells in active drug-treated wells, and by using histograms produced by Columbus to visualize gates applied to the distribution of measured/calculated values for each feature.

### Normalized growth rate inhibition (GR) metrics

Viable cell counts at time=0 and time=72h as determined in Columbus were then exported and used to compute GR values and GR metrics with the freely available online GR calculator (http://www.grcalculator.org/grcalculator/), which is based on previously published methods (Hafner et al., 2016). Dose response curves of GR values were produced using MATLAB (MathWorks).

### Quantification of immunofluorescence images

Immunofluorescence intensities in the entire cell or relevant subcellular compartment were measured in Columbus. Masks for the nucleus, cytoplasm and entire cell were generated by thresholding the Hoechst and the Whole Cell Stain channels. Custom masks were generated for each cell line due to variation in cell morphology. To quantify changes in the levels of proteins/ phosphoproteins after treatment with DMSO or active drug, the relevant mask (as determined both by prior knowledge and by the observed staining pattern for each antibody) was applied to microscopy images and the fluorescence intensity was measured in that specific region of each cell. Levels of phospho-AKT (T308 and S473), phospho-4E-BP1 and phospho-GSK3β in single cells were measured using a whole-cell mask. Phospho-S6 levels were measured within a cytoplasmic mask. Signals for BrdU, EdU, F-ara-EdU, cyclin A2, cyclin D1, FOXO3, geminin, p21, p27, phospho-Chk1, phospho-Chk2, phospho-H2A.X, phospho-Histone H3, phospho-pRb and phospho-RPA levels were measured within a nuclear mask. Total DNA content and EdU content per nucleus were calculated by multiplying the average nuclear fluorescence intensity by the nuclear area. The PI3K signaling index was computed by adding the mean values (normalized to DMSO) of each of nine different proteins/phosphoproteins (scale 0-9, with 9 indicating no difference from DMSO). For signals that increase with PI3K pathway suppression (nuclear FOXO3, nuclear p21, and nuclear p27), the inverse value was used when computing the index. “Percent high” for any measured protein/post-translational modification was determined by applying an identical threshold to the entire distribution of values determined for cells treated with DMSO or active drug (see **Figure 4A**). All raw data were exported from Columbus; visualization, plotting and statistical analysis of the data were performed in MATLAB. Specific gating strategies are illustrated in individual figure panels; experimental replicates and statistical analyses are described in figure legends. Cells with mitotic aberrations were quantified based on visual inspection of individual microscopy images.

### Cell cycle analysis and cell cycle stage-specific immunofluorescence values

After masking the nuclei in Columbus, average Hoechst and EdU intensity values were extracted and multiplied by the nuclear area to determine the DNA content and EdU content, respectively, on a single-cell level basis. The bivariate plot of nuclear DNA content versus EdU content readily separated cells into three groups representing G1, S and G2/M cells (see **Figure S3A**). S-phase-non-replicating (SNR) cells were identified based on having DNA content intermediate to G1 and G2/M and no EdU signal. The percentages of cells in different cell cycle stages were determined in MATLAB by gating different groups of cells by DNA content (X) and EdU content (Y). In each experiment, the specific values for gates were first determined using data for cells treated with DMSO only, and then applied to data for drug-treated cells. For analysis of immunofluorescence intensities in specific subpopulations of cells defined by cell cycle stage (e.g., **Figure 2D**), nuclear intensity values from cells labeled with Hoechst/EdU and immunostained for a nuclear marker of interest were first plotted in MATLAB using the 3-way scatterplot function to preserve relationships among the three measured intensity values in single cells. Cells in different cell cycle stages were then gated as described above. The entire distribution of immunofluorescence intensity values for cells in each cell cycle stage was then replotted in relation to DNA content.

### Live-cell analysis of proliferation/apoptosis

To determine population growth trajectories, total cell number in an entire well imaged using the 4x lens was determined using Incucyte Zoom software (Essen Bioscience). Cell number was calculated by identifying nuclei positive for GFP (HCC70) or mCherry (Hs 578T, HCC1806) in microscopy images, based on a background-subtracted fluorescence intensity threshold. C3/7-positive cells were similarly identified based on a fluorescence intensity threshold in the GFP channel. For each of three technical replicates in each independent experiment, C3/7 counts over time were plotted and area under the curve (AUC) calculations were determined in MATLAB and averaged. The mean fold-change value was then calculated as the ratio of the average AUC for drug versus the average for DMSO. Mean fold-change values were then averaged across independent experiments.

### Live-cell analysis of drug effect on the cell cycle

Semi-automated tracking of single HCC1806 cells expressing H2B-mTurquoise, mVenus-hGeminin(1-110) and mCherry-dE2F1N was performed using p53CinemaManual 2.0, a MATLAB-based software package freely available at: https://github.com/balvahal/p53CinemaManual/releases/tag/v2.0.0. Individual cells were tracked in time-lapse movies using H2B-mTurquoise; fluorescence intensities for mVenus and mCherry were then extracted from each tracked cell by the software. Despite FACS sorting, many cells were not double-positive for mVenus and mCherry. Thus, cell cycle analysis was performed by analyzing plots of mVenus intensity alone versus time (see annotated trace of representative single cell in DMSO, **Figure 3A**). Each individual trace was manually analyzed to ascertain the timing of cell cycle stage transitions. The data for each cell were then compiled into matrix that underlies each heatmap. In the mTurquoise channel, cell divisions were scored by visualizing chromosome segregation followed by appearance of daughter cell nuclei. Cell death was recorded by the appearance of nuclear fragmentation (karyorrhexis).

### Analysis of steady-state polar metabolites

Metabolite peak intensities for drug-treated samples were compared to peak intensities for samples treated with DMSO (N=3 independent experiments) using MetaboAnalyst 4.0 (MA) (https://www.metaboanalyst.ca/MetaboAnalyst/faces/home.xhtml). Prior to uploading the data, an initial processing step was performed to exclude metabolites with two or more missing values in either the DMSO or active treatment group. For rapamycin, AZD8055, and Torin2, 34 (11.2%), 35 (11.6%), and 34 (11.2%) of 303 metabolites, respectively, were excluded for this reason. For a small number of excluded metabolites (4/34 for rapamycin, 8/35 for AZD8055, and 9/34 for Torin2), three numerical values were available for DMSO but only 0 or 1 for active treatment, suggesting that treated samples had undetectable levels by mass spectrometry due to low abundance. Because the fold-change values for these metabolites were incalculable but in certain cases could be large, these metabolites and their raw values are listed in separate tables in **Table S4**. To facilitate downstream analysis in MA, single missing values in either treatment group were imputed by taking the average of the two measured values. This step was performed prior to analyzing the data in MA because the software performs missing value estimation for each metabolite using all of the sample data rather than the data for individual treatment groups. In certain instances, this method was observed to obscure strong changes in metabolite levels induced by drug treatment. Once the data were uploaded to MA, no additional filtering was applied. Sample data were quantile-normalized, and then the metabolite data were log-transformed. No scaling was performed. The effects of data normalization and transformation were visualized using diagnostic plots. Fold-change values were calculated as the ratio between group means, using absolute values of metabolite data prior to log-transformation. T-tests were performed to assess the statistical significance of fold-change values. Volcano plots were generated using fold-change values and p-values from T-tests to identify metabolites with the largest and most significant changes after drug treatment.

### DATA AND SOFTWARE AVAILABILITY

### Datasets being prepared for release

1. Evaluation of the sensitivity of six TNBC cell lines to 23 PI3K pathway drugs or trametinib (MEK inhibitor). Data summarized in **Figures 1B**, **S1D**.

a. Measured GR values (72h)
b. Calculated GR metrics (72h)
2. Evaluation of the sensitivity of HCC1806 and HCC70 cells to Torin1, Torin2, AZD8055 or nine Torin2 analogs. Data summarized in **Figures 5B-C**.

a. Measured GR values (72h)
b. Calculated GR metrics (72h)
3. Targeted metabolomics profiling data (i.e., peak intensity measurements) of BT-549 cells after exposure of to DMSO, rapamycin (1μM), AZD8055 (1μM), or Torin2 (1μM) for 6 hours. Data summarized in **Figure 2F**.

### Datasets currently available

1. Evaluation of the sensitivity of Torin2 compared to the PI3K pathway drugs alpelisib, TGX-221, pictilisib, everolimus, and sapanisertib in a collection of breast cancer cell lines (N=28). Data summarized in **Figures 1C, S1E.**

a. Measured GR values (72h): http://lincs.hms.harvard.edu/db/datasets/20343/
b. Calculated GR metrics (72h): http://lincs.hms.harvard.edu/db/datasets/20344/

