## Supplementary Figures and Tables for "Combined inhibition of mTOR and PIKKs exploits replicative and checkpoint vulnerabilities to induce death of PI3K-activated triple-negative breast cancer cells"

#### SUPPLEMENTARY INFORMATION

Seven supplementary figures with legends below.

#### SUPPLEMENTARY FIGURE LEGENDS

**Figure S1 (related to Figure 1: Torin2 and omipalisib produce strong cytotoxic and anti-proliferative effects in TNBC cells)**. **A.** Quantitative western blotting of PTEN and INPP4B protein levels in 46 breast cancer and non-transformed mammary epithelial cell lines (see **Table S1**). The plot shows 48 data points because two independent protein lysates were prepared from SK-BR-3 cells and because MDA-MB-231 cells were cultured in two different types of media. Shown are mean values from at least 2 independent experiments. TNBC/basal-like cell lines are marked in red. Six TNBC/basal-like cell lines selected for more focused study are numbered. **B.** Cell counts at the beginning and end of the 72h treatment period were determined by co-staining with Hoechst (blue) and Live/Dead cell viability stain (red), followed by microscopy and image analysis (see STAR methods for details). Shown are representative images of HCC1806 cells following treatment with $GR_{max}$ doses (3.2μM) of AZD8055 or Torin2 for 72h. Dead cells were identified based on pyknosis/karyorrhexis in the Hoechst channel and positive Live/Dead staining in the far red channel and excluded. Viable cell counts were then used to produce nine-point dose response curves, and GR metrics were computed to determine relative potency and efficacy. **C.** GR metrics quantify variation in drug potency and maximal efficacy. $GR_{max}$ >0 indicates partial growth inhibition, $GR_{max}$ =0 indicates complete growth inhibition, and $GR_{max}$ <0 indicates cytotoxicity. **D.** Clustergram heatmap of mean $GR_{50}$ values for 24 drugs in six TNBC cell lines with PI3K pathway dysregulation; N=3 experiments, each performed in quadruplicate. The three drugs with the lowest $GR_{max}$ values (**Figure 1B**) are highlighted in red. Trametinib (blue) is a MAPK pathway inhibitor included as a comparator. See **Table S3** for details on all drugs. **E.** $GR_{max}$ values for a subset of PI3K pathway drugs including Torin2 in 7 luminal and 2 non-transformed mammary epithelial cell lines. *P<0.05, **P<0.01, ***P<0.001, and n.s. (not significant) by Mann-Whitney U test. **F.** Growth curves for HCC1806 NLS-mCherry cells in the presence of $GR_{max}$ concentrations of AZD8055, omipalisib and Torin2 (all 3.2μM) or 0.1x$GR_{max}$ concentrations of Torin2 and omipalisib (0.32μM). Data points/shading indicate mean ± SD of 3 replicates in a single time-lapse experiment; N=3 experiments, representative data shown. "*" denotes start of treatment. **G.** Percentage of unlabeled nuclei after 48 or 75h of continuous labeling of HCC1806 cells with F-ara-EdU, starting after 24h of treatment. Drugs were used at $GR_{max}$ concentrations (3.2μM) except for Torin2 and omipalisib, which were used at 0.3x$GR_{max}$ (1μM) due to cell death at 3.2μM. Shown are the mean ± SEM; N=3 experiments (see **Figure 1E** for representative data from a single experiment). **H.** Representative images of Hs 578T cells in the presence or absence of a fluorescent reporter of caspase 3/7 activity. Cells were exposed to $GR_{max}$ concentrations of AZD8055 (1μM) or Torin2 (3.2μM) for 24h, as indicated. Images are shown in greyscale for clarity. "BF" indicates brightfield image. Scale bar=400μm.

**Figure S2 (related to Figure 2: PI3K/AKT/mTOR inhibitors impede progression of S phase.). A**. p-AKT S473 levels in HCC1806 cells by immunofluorescence after 24h of drug exposure. Scale bar=50μm. **B.** Levels of p-AKT S473 in single cells. Boxplots show the median and 25th/75th percentiles. **C.** Mean ± SEM levels of six proteins/phosphoproteins whose values are expected to decrease with inhibition of PI3K pathway activity; N≥2 experiments, each performed in triplicate. Cell lines are as indicated. Protein/phosphoprotein levels were used to generate the clustergram heat maps shown in **Figure 2A** and calculate the signaling index values shown in **Figure S2D**. Measurements were performed by quantitative immunofluorescence microscopy after 24h of exposure to $GR_{max}$ concentrations (1-3.2μM) of the indicated drugs. In HCC1806 cells only, Torin2 and omipalisib were used at 0.3x$GR_{max}$ (1μM) due to cell death. Fluorescence intensity values were extracted from whole cells (p-AKT T308, p-AKT S473, p-GSK3β S9, p-4E-BP1 T37/46) or cytoplasmic (p-S6 S235/236) or nuclear (cyclin D, FOXO3, p21 Cip1, p27 Kip1) compartments by image analysis. For each protein/phosphoprotein, background-subtracted fluorescence intensity measurements were normalized to values for DMSO. **D**. Mean ± SEM of signaling index values for the data shown in C. Index values range from 0 for complete inhibition to 9 for no difference from DMSO-only control (see STAR methods for details). **E.** Mean percent ± SEM of p-pRb-low HCC1806 cells t=$T_d$ (28h); N=3 experiments. ***P<0.001 by one-way ANOVA and Tukey's test (only selected comparisons shown).

**Figure S3 (related to Figure 2: PI3K/AKT/mTOR inhibitors impede progression of S phase.). A**. Analysis of nuclear DNA content vs EdU content in BT-20 cells treated with 3.2μM alpelisib or DMSO. Colors indicate cell cycle stage: G1 (black), active S phase (red), S-phase non-replicating ($S_{NR}$; yellow), and G2/M (blue). Color scheme applies to data in panels B, E, F and H. **B.** Distribution of cells in different cell cycle stages in two additional TNBC/basal-like cell lines after exposure to the indicated treatments for t= $T_d$ (32h for Hs 578T, 40h for BT-549). Drugs were used at $GR_{max}$ doses (1-3.2μM) except for Torin2 and omipalisib, which were used at 0.3x$GR_{max}$ concentrations (1μM) in Hs 578T cells due to cell death. Drugs are ordered by viable cell number at $T_d$, with the least effective on top and most effective on the bottom. **C.** DNA content vs mean nuclear EdU intensity after exposure of HCC1806 cells to Torin2 at 0.03-0.3x$GR_{max}$ dose. DNA content distribution for gated EdU-negative cells (red box) is shown below. Black arrows denote $S_{NR}$ cells. **D.** Mean ± SEM of active S vs. $S_{NR}$ cells from data shown in C; N=3 experiments. **E**. DNA content vs. mean nuclear intensity values for p-pRb after exposure of HCC1806 cells to $GR_{max}$ doses of AZD8055 or Torin2 for t=$T_d$. Colors indicate different cell cycle stages based on measurement of DNA content and EdU content in the same cells. **F.** Effects of submaximal concentrations (0.1-1x$GR_{max}$) of AZD8055 on the distribution of BT-549 cells in different cell cycle stages. Submaximal doses were used to analyze S-phase cells because few cells were in active S phase at $GR_{max}$ doses (1μM). Cells were pulse-labeled with EdU for 60min. **G.** Bar graphs (left) display the percentage of EdU-positive (active) S-phase cells in the entire cell population after treatment with DMSO or AZD8055 at the indicated doses. Boxplots overlaid onto single cell data (right) indicate the median and $25^{th}/75^{th}$ percentiles for nuclear EdU content in all S-phase cells (active S plus $S_{NR}$). **H.** Cell cycle analysis for BT-20 cells performed after 6h of exposure to the indicated drugs at $GR_{max}$ concentrations (1-3.2μM). **I.** Quantification of nuclear EdU content in S-phase cells after 6h (0.125 x $T_d$) or 14h (0.3 x $T_d$) of drug exposure. ***P<0.001 vs DMSO control (Mann-Whitney U test).

**Figure S4 (related to Figure 4: Torin2 causes replication catastrophe). A.** Torin2 (but not omipalisib) increases the percentage of HCC70 cells with high levels of native BrdU, γH2A.X, and p-RPA at t= ~0.5x$T_d$ (20-24h). Cells with increased nuclear levels appear in the red gate and are labeled red. Drugs were used at $GR_{max}$ doses (1-3.2μM) except for Torin2, which was used at 0.3x$GR_{max}$ (1μM). **B.** Bar graphs display the mean ± SEM; N=3 experiments. *P<0.05, **P<0.01, ***P<0.001 vs. DMSO by one-way ANOVA and Dunnett's test. **C.** DNA content vs. nuclear intensities of p-Chk1 S317 and p-Chk2 T68 after treating HCC1806 cells with the indicated drugs for ~0.2x$T_d$ (5h). Cells with increased nuclear levels appear in the red gate and are labeled red. Quantification of independent experimental replicates appears in **Figure 4C**.

**Figure S5 (related to Figure 5: The activity of Torin2 in TNBC results from combined antagonism of mTORC1/2 and PIKKs). A.** Dose-response curves (72h) for AZD8055, Torin2, AZ20, and AZD8055/AZ20 mixed at equimolar concentrations in HCC1806 and HCC70 cells. **B.** Chemical structures of Torin1, Torin2 and nine Torin2 analogs. **C.** Levels of p-AKT S473, p-4E-BP1 T37/46 and γH2A.X at t=0.33x$T_d$ (14h) vs. GR values computed after 72h of exposure of HCC70 cells to 0.032, 0.32, or 3.2μM of AZD8055, Torin1, Torin2 or Torin2 analogs. Drugs/doses whose GR values correlate with mTORC1/2 signaling activity fall on the black regression lines (*left, middle*). Drugs/doses whose activities are discontinuous with the regression fall within the red boxes (*left, middle*) and cause increased levels of γH2A.X (*black box*, *right*). **D.** DNA content vs. nuclear γH2A.X or p-pRb levels at 14h for DMSO (#1) and drugs/doses whose activities correlate with mTORC1/2 signaling activity (#2-4) and those that do not (#5-8). Percentages of gated γH2A.X-high or p-pRb-low cells are relative to entire treated population. **E**. Mean percent ± SEM of γH2A.X-high cells or p-pRb-low cells at 14h post-treatment, based on data shown in D; N=3 experiments. *P<0.05, **P<0.01,***P<0.001 by one-way ANOVA and Dunnett's test.

**Figure S6 (related to Figure 6: Combined inhibition of mTORC1/2 and PIKKs produces benefit by co-targeting pathways required in S phase). A.** Results of analysis of interactions between PI3K pathway drugs and the Chk1 inhibitor rabusertib. "+" indicates additive or greater benefit by Loewe criteria, "N.I." indicates no interaction, and "ant." indicates antagonism. **B.** Dose ranges of drugs used for analysis of drug interactions. Ranges were selected based on the dose response curves for each single agent and reflect submaximal levels of drug effect (spanning the $GR_{50}$ dose). **C.** Analysis of mTORC1/2 signaling after exposure of HCC70 or Hs 578T cells to 32-100nM of AZD8055, which is equivalent to 0.01-0.03x$GR_{max}$ dose in HCC70 cells and 0.03-0.1x$GR_{max}$ dose in Hs 578T cells. HCC70 and Hs 578T cells were exposed to drugs for t= ~0.5x$T_d$ (24h) and for t=0.75x$T_d$ (24h), respectively. The effects of 1μM Torin2 are shown for comparison.

**Figure S7 (related to Figure 7. Low doses of mTORC1/2 and ATR/Chk1 inhibitors in combination cause increased ssDNA in mitotic prophase and death). A.** Mitoses per high-powered field (h.p.f.) after exposure of HCC70 cells to the indicated drugs for t= ~0.5x$T_d$ (24h). Mitoses were quantified by staining cells with Hoechst and anti-CREST, anti-β-tubulin, and anti-γ-tubulin antibodies. Bar graphs represent the mean ± SEM; N=3 experiments. Values were normalized to DMSO prior to averaging.

*P<0.05, **P<0.01, ***P<0.001 and n.s. (not significant) by ANOVA and Tukey's test (only selected comparisons shown). **B.** Mean percent ± SEM of cells in different stages of mitosis, as determined by immunofluorescence microscopy; N=3 experiments, 569 mitoses. *P<0.05, **P<0.01, ***P<0.001 and n.s. (not significant) by ANOVA and Tukey's test (only selected comparisons shown). **C.** Rate of mitotic entry (% of cells per hour) as determined by time-lapse imaging of asynchronous HCC70 H2B-GFP cells after 15 or 30h of exposure to the indicated treatments.

### FIGURE S1

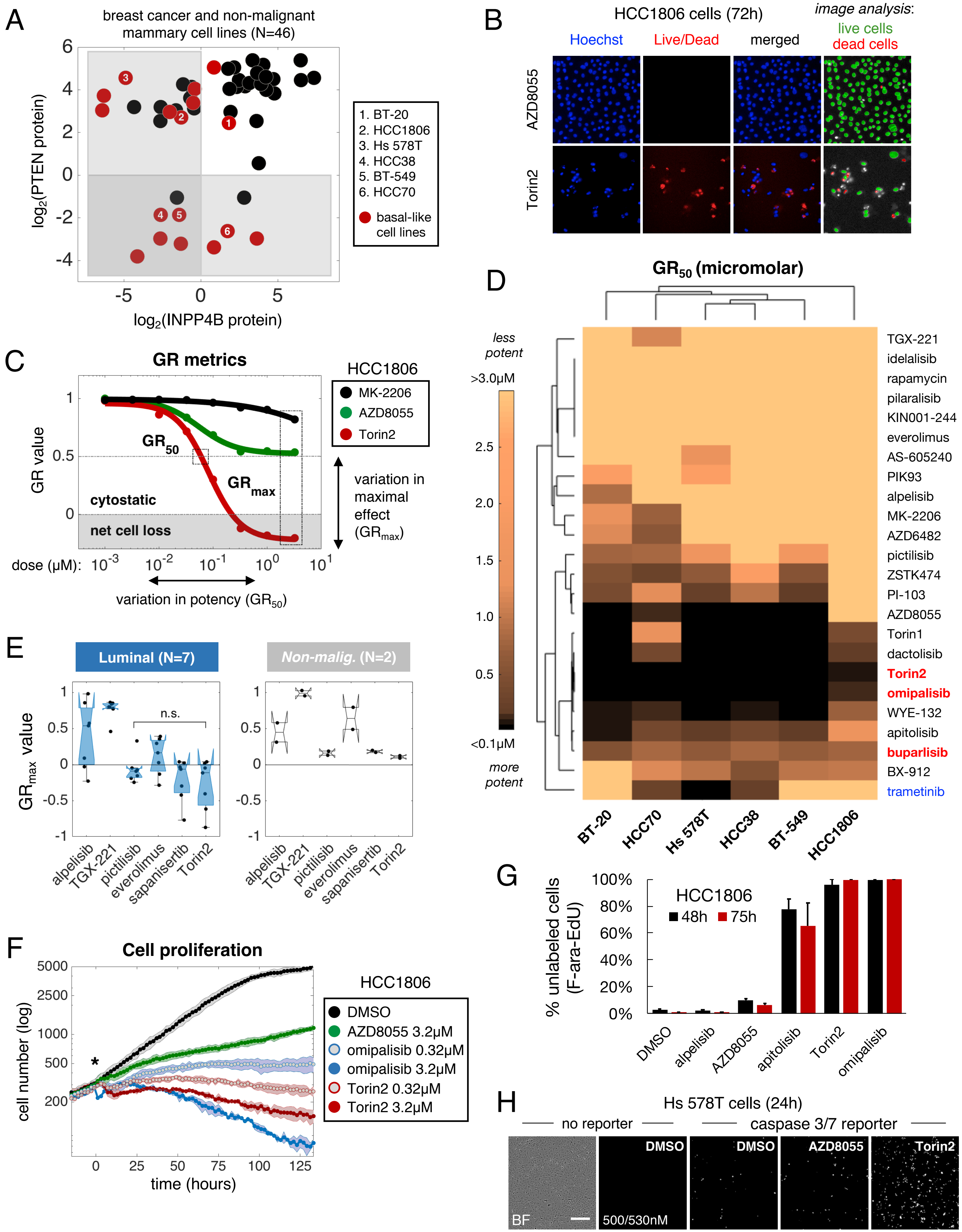
A
breast cancer and non-malignant mammary cell lines (N=46)
log2(PTEN protein)
log2(INPP4B protein)
1. BT-20
2. HCC1806
3. Hs 578T
4. HCC38
5. BT-549
6. HCC70
basal-like cell lines
B
HCC1806 cells (72h)
Hoechst
Live/Dead
merged
image analysis:
live cells
dead cells
AZD8055
Torin2
C
GR metrics
HCC1806
MK-2206
AZD8055
Torin2
GR value
GR50
GRmax
cytostatic
net cell loss
variation in maximal effect (GRmax)
dose (µM):
variation in potency (GR50)
D
GR50 (micromolar)
less potent
>3.0µM
<0.1µM
more potent
TGX-221
idelalisib
rapamycin
pilaralisib
KIN001-244
everolimus
AS-605240
PIK93
alpelisib
MK-2206
AZD6482
pictilisib
ZSTK474
PI-103
AZD8055
Torin1
dactolisib
Torin2
omipalisib
WYE-132
apitolisib
buparlisib
BX-912
trametinib
BT-20
HCC70
Hs 578T
HCC38
BT-549
HCC1806
E
Luminal (N=7)
Non-malig. (N=2)
GRmax value
n.s.
alpelisib
TGX-221
pictilisib
everolimus
sapanisertib
Torin2
F
Cell proliferation
cell number (log)
time (hours)
HCC1806
DMSO
AZD8055 3.2µM
omipalisib 0.32µM
omipalisib 3.2µM
Torin2 0.32µM
Torin2 3.2µM
G
% unlabeled cells (F-ara-EdU)
HCC1806
48h
75h
DMSO
alpelisib
AZD8055
apitolisib
Torin2
omipalisib
H
Hs 578T cells (24h)
no reporter
caspase 3/7 reporter
BF
DMSO
500/530nM
DMSO
AZD8055
Torin2

FIGURE S2

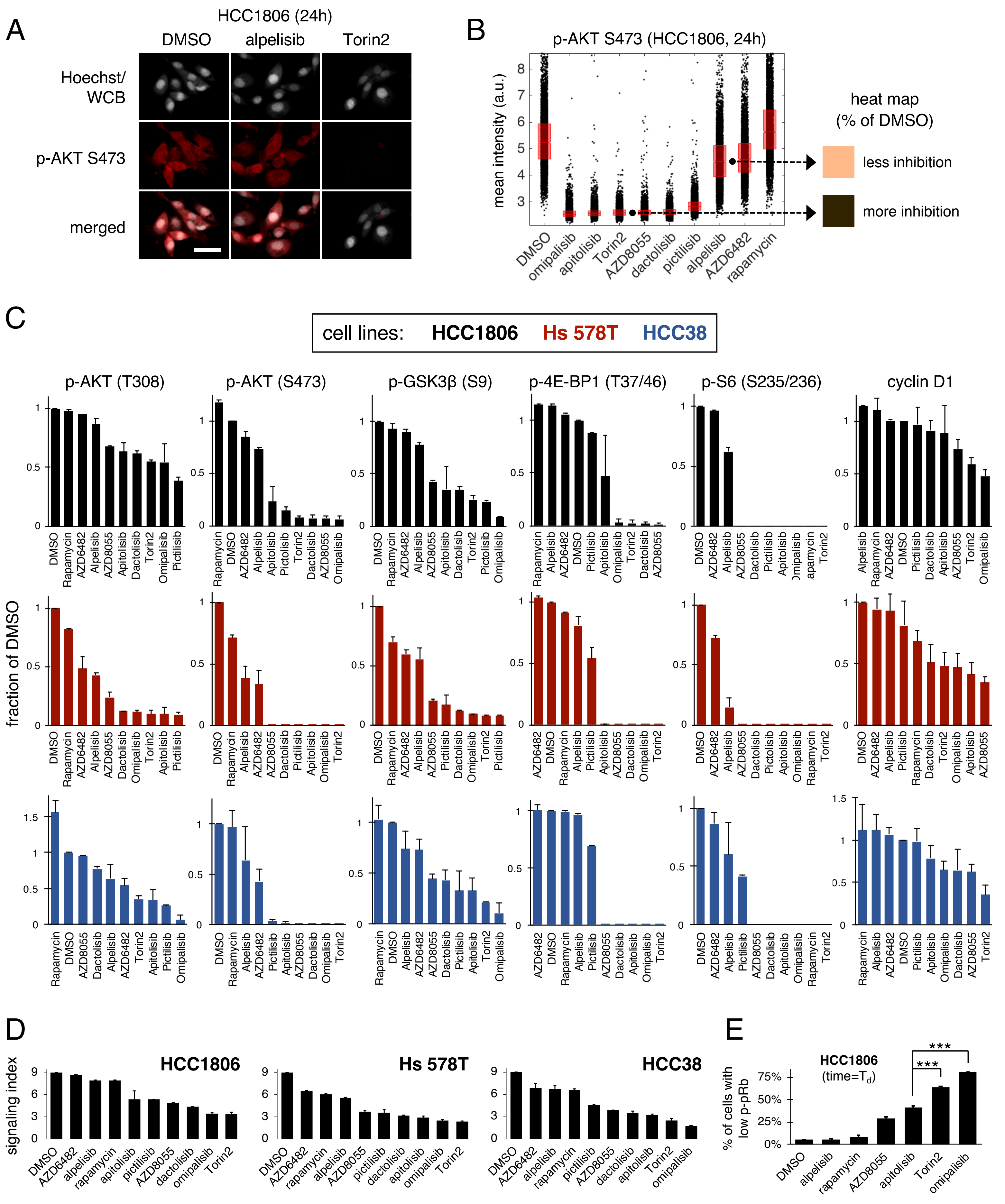
A
HCC1806 (24h)
DMSO
alpelisib
Torin2
Hoechst/
WCB
p-AKT S473
merged
B
p-AKT S473 (HCC1806, 24h)
mean intensity (a.u.)
DMSO
omipalisib
apitolisib
Torin2
AZD8055
dactolisib
pictilisib
alpelisib
AZD6482
rapamycin
heat map
(% of DMSO)
less inhibition
more inhibition
C
cell lines:
HCC1806
Hs 578T
HCC38
p-AKT (T308)
p-AKT (S473)
p-GSK3β (S9)
p-4E-BP1 (T37/46)
p-S6 (S235/236)
cyclin D1
fraction of DMSO
DMSO
Rapamycin
AZD6482
Alpelisib
AZD8055
Apitolisib
Dactolisib
Torin2
Omipalisib
Pictilisib
D
signaling index
HCC1806
Hs 578T
HCC38
E
HCC1806
(time=T$_d$)
% of cells with
low p-pRb
***
***
DMSO
alpelisib
rapamycin
AZD8055
apitolisib
Torin2
omipalisib

### FIGURE S3

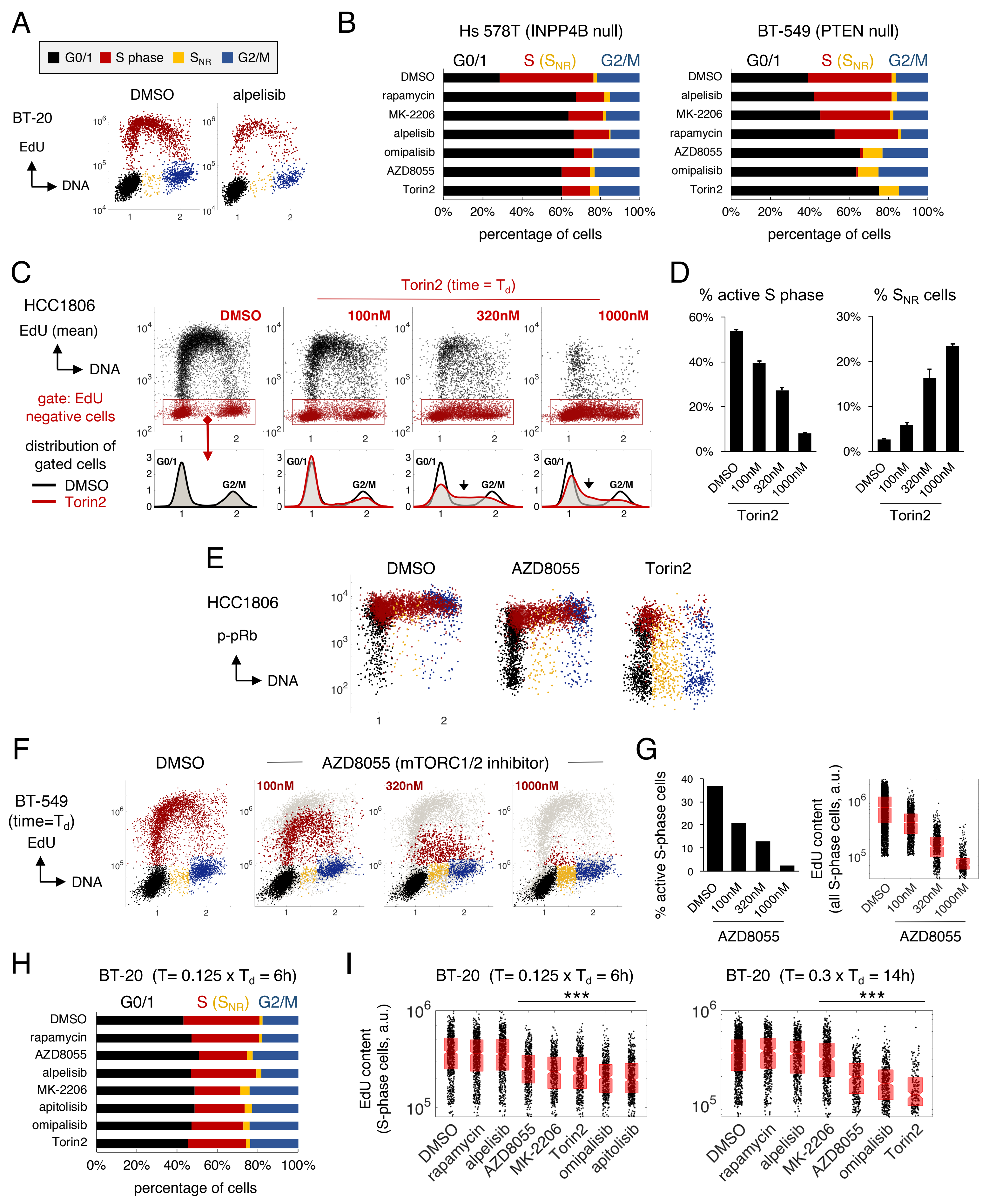

A
G0/1 S phase S_NR G2/M
DMSO
alpelisib
BT-20
EdU
DNA
B
Hs 578T (INPP4B null)
BT-549 (PTEN null)
G0/1 S (S_NR) G2/M
DMSO
rapamycin
MK-2206
alpelisib
omipalisib
AZD8055
Torin2
percentage of cells
C
HCC1806
EdU (mean)
DNA
gate: EdU negative cells
distribution of gated cells
DMSO
Torin2
Torin2 (time = T_d)
100nM
320nM
1000nM
G0/1
G2/M
D
% active S phase
% S_NR cells
Torin2
E
HCC1806
p-pRb
DNA
DMSO
AZD8055
Torin2
F
BT-549 (time=T_d)
EdU
DNA
DMSO
AZD8055 (mTORC1/2 inhibitor)
100nM
320nM
1000nM
G
% active S-phase cells
EdU content (all S-phase cells, a.u.)
AZD8055
H
BT-20 (T= 0.125 x T_d = 6h)
G0/1 S (S_NR) G2/M
DMSO
rapamycin
AZD8055
alpelisib
MK-2206
apitolisib
omipalisib
Torin2
percentage of cells
I
BT-20 (T= 0.125 x T_d = 6h)
BT-20 (T= 0.3 x T_d = 14h)
EdU content (S-phase cells, a.u.)
***
DMSO
rapamycin
alpelisib
AZD8055
MK-2206
Torin2
omipalisib
apitolisib

### FIGURE S4

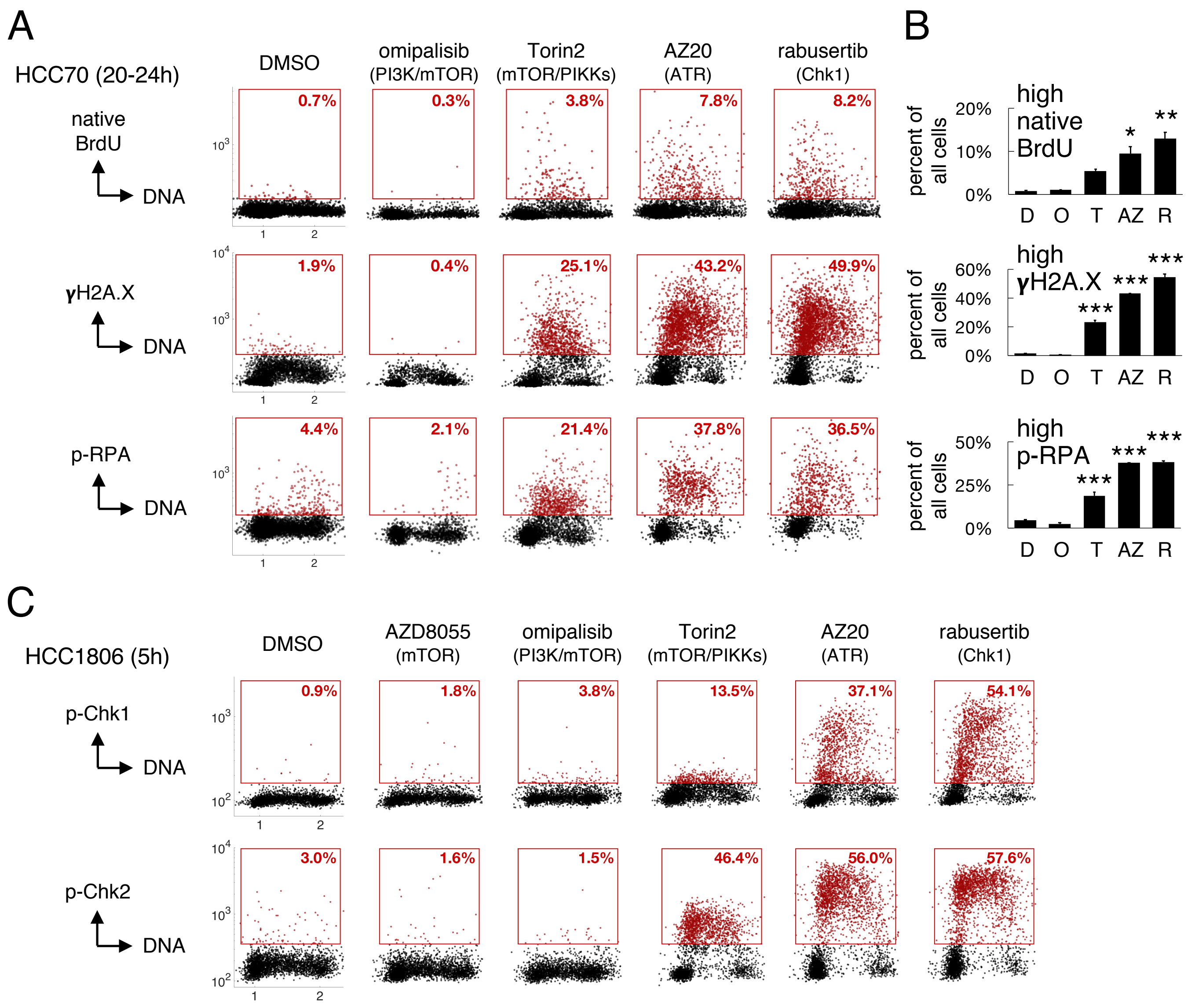
A
HCC70 (20-24h)
DMSO
omipalisib
(PI3K/mTOR)
Torin2
(mTOR/PIKKs)
AZ20
(ATR)
rabusertib
(Chk1)
native
BrdU
DNA
0.7%
0.3%
3.8%
7.8%
8.2%
γH2A.X
DNA
1.9%
0.4%
25.1%
43.2%
49.9%
p-RPA
DNA
4.4%
2.1%
21.4%
37.8%
36.5%
B
percent of
all cells
high
native
BrdU
20%
10%
0%
D O T AZ R
high
γH2A.X
60%
40%
20%
0%
high
p-RPA
50%
25%
0%
C
HCC1806 (5h)
DMSO
AZD8055
(mTOR)
omipalisib
(PI3K/mTOR)
Torin2
(mTOR/PIKKs)
AZ20
(ATR)
rabusertib
(Chk1)
p-Chk1
DNA
0.9%
1.8%
3.8%
13.5%
37.1%
54.1%
p-Chk2
DNA
3.0%
1.6%
1.5%
46.4%
56.0%
57.6%

FIGURE S5

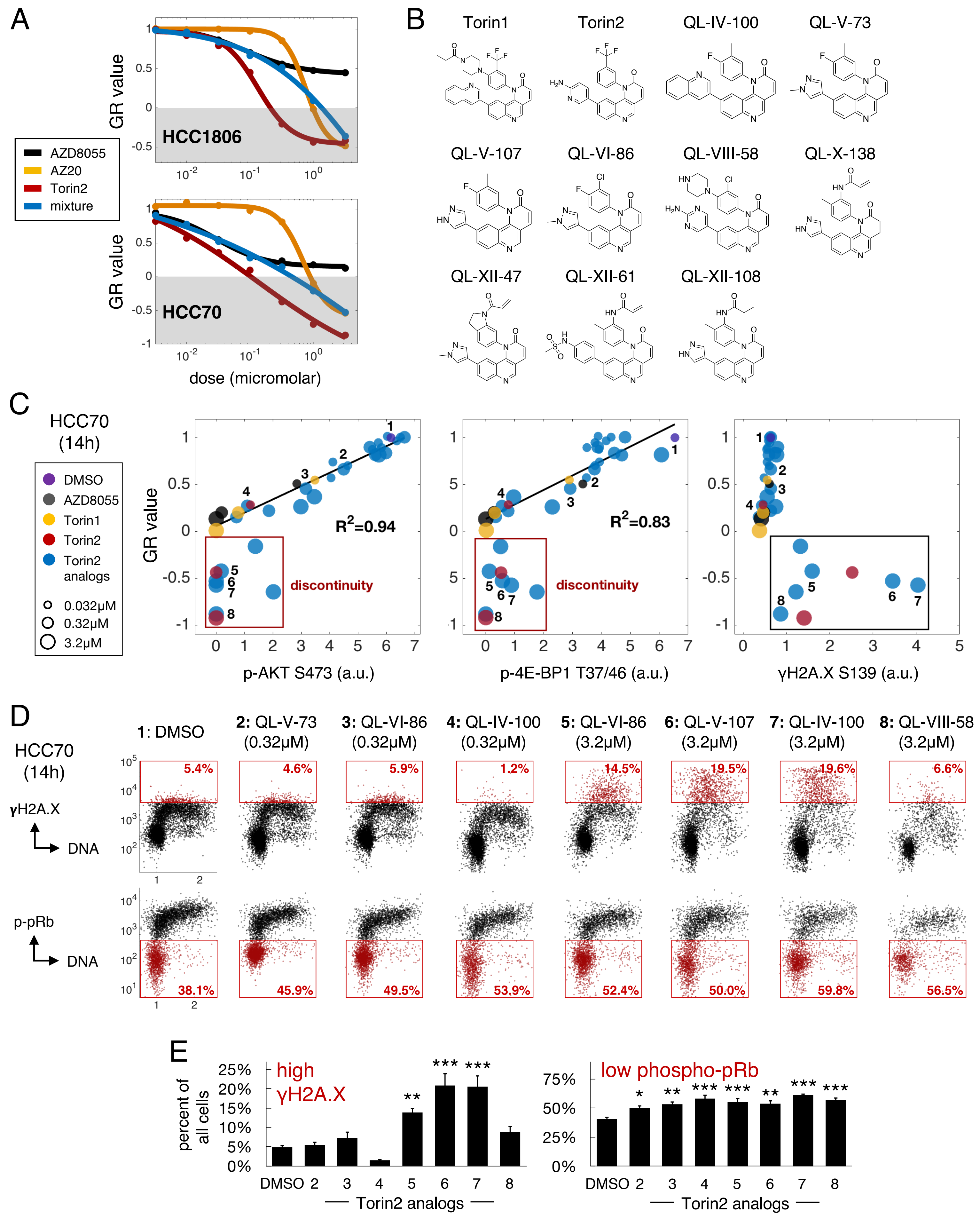

A
GR value
HCC1806
HCC70
AZD8055
AZ20
Torin2
mixture
dose (micromolar)
B
Torin1
Torin2
QL-IV-100
QL-V-73
QL-V-107
QL-VI-86
QL-VIII-58
QL-X-138
QL-XII-47
QL-XII-61
QL-XII-108
C
HCC70 (14h)
DMSO
AZD8055
Torin1
Torin2
Torin2 analogs
0.032μM
0.32μM
3.2μM
GR value
R2=0.94
R2=0.83
discontinuity
p-AKT S473 (a.u.)
p-4E-BP1 T37/46 (a.u.)
γH2A.X S139 (a.u.)
D
HCC70 (14h)
γH2A.X
DNA
p-pRb
DNA
1: DMSO
2: QL-V-73 (0.32μM)
3: QL-VI-86 (0.32μM)
4: QL-IV-100 (0.32μM)
5: QL-VI-86 (3.2μM)
6: QL-V-107 (3.2μM)
7: QL-IV-100 (3.2μM)
8: QL-VIII-58 (3.2μM)
5.4%
4.6%
5.9%
1.2%
14.5%
19.5%
19.6%
6.6%
38.1%
45.9%
49.5%
53.9%
52.4%
50.0%
59.8%
56.5%
E
percent of all cells
high γH2A.X
low phospho-pRb
DMSO 2 3 4 5 6 7 8
Torin2 analogs

### FIGURE S6

A

| | rabusertib (Chk1i) | | | | |
|---|---|---|---|---|---|
| | HCC70 | Hs 578T | BT-549 | HCC1806 | BT-20 |
| AZD6482 | NI | NI | NI | NI | + |
| AZD8055 | + | + | + | + | + |
| buparlisib | NI | NI | NI | NI | + |
| alpelisib | NI | NI | NI | NI | + |
| apitolisib | + | + | + | + | + |
| MK-2206 | + | NI | NI | NI | + |
| rapamycin | NI | NI | + | NI | + |
| WYE-132 | + | + | + | + | + |

B

dose ranges evaluated

| | **HCC70** | **Hs 578T** | **BT-549** | **HCC1806** | **BT-20** |
|---|---|---|---|---|---|
| **AZ20** | 100-1000 | 320-3200 | 320-3200 | 100-1000 | 320-3200 |
| **rabusertib** | 100-1000 | 100-1000 | 320-3200 | 60-600 | 1000-5600 |
| AZD6482 | 3.2-100 | 320-3200 | 320-3200 | 100-1000 | 320-3200 |
| AZD8055 | 10-200 | 10-200 | 10-200 | 10-1000 | 10-320 |
| buparlisib | 100-1000 | 100-1000 | 100-1000 | 100-1000 | 100-1000 |
| alpelisib | 320-3200 | 320-3200 | 320-3200 | 320-3200 | 320-3200 |
| apitolisib | 3.2-320 | 32-320 | 32-2000 | 100-3200 | 32-320 |
| MK-2206 | 100-1000 | 320-3200 | 100-3200 | 320-3200 | 320-3200 |
| rapamycin | 100-1000 | 100-1000 | 100-1000 | 100-1000 | 100-1000 |
| WYE-125 | 10-200 | 10-200 | 10-400 | 10-1000 | 10-320 |

C

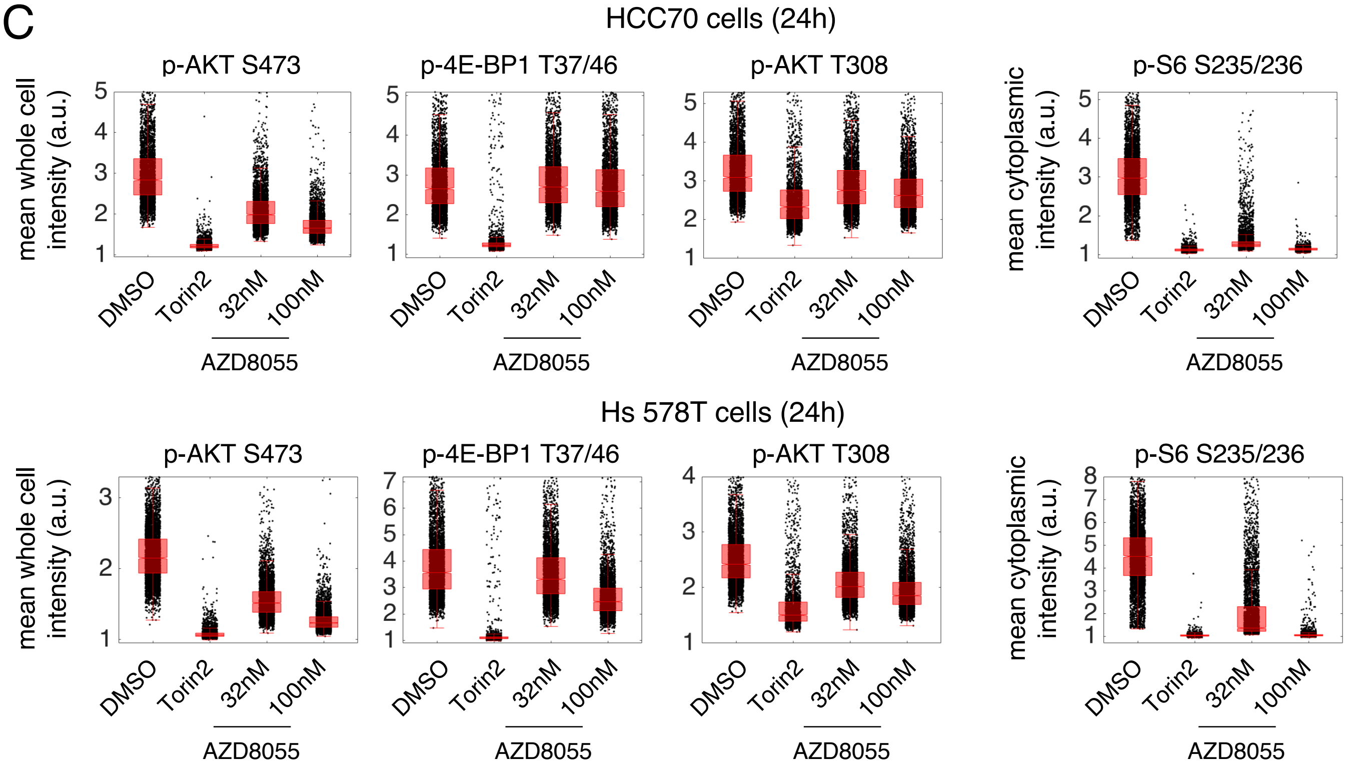

### FIGURE S7

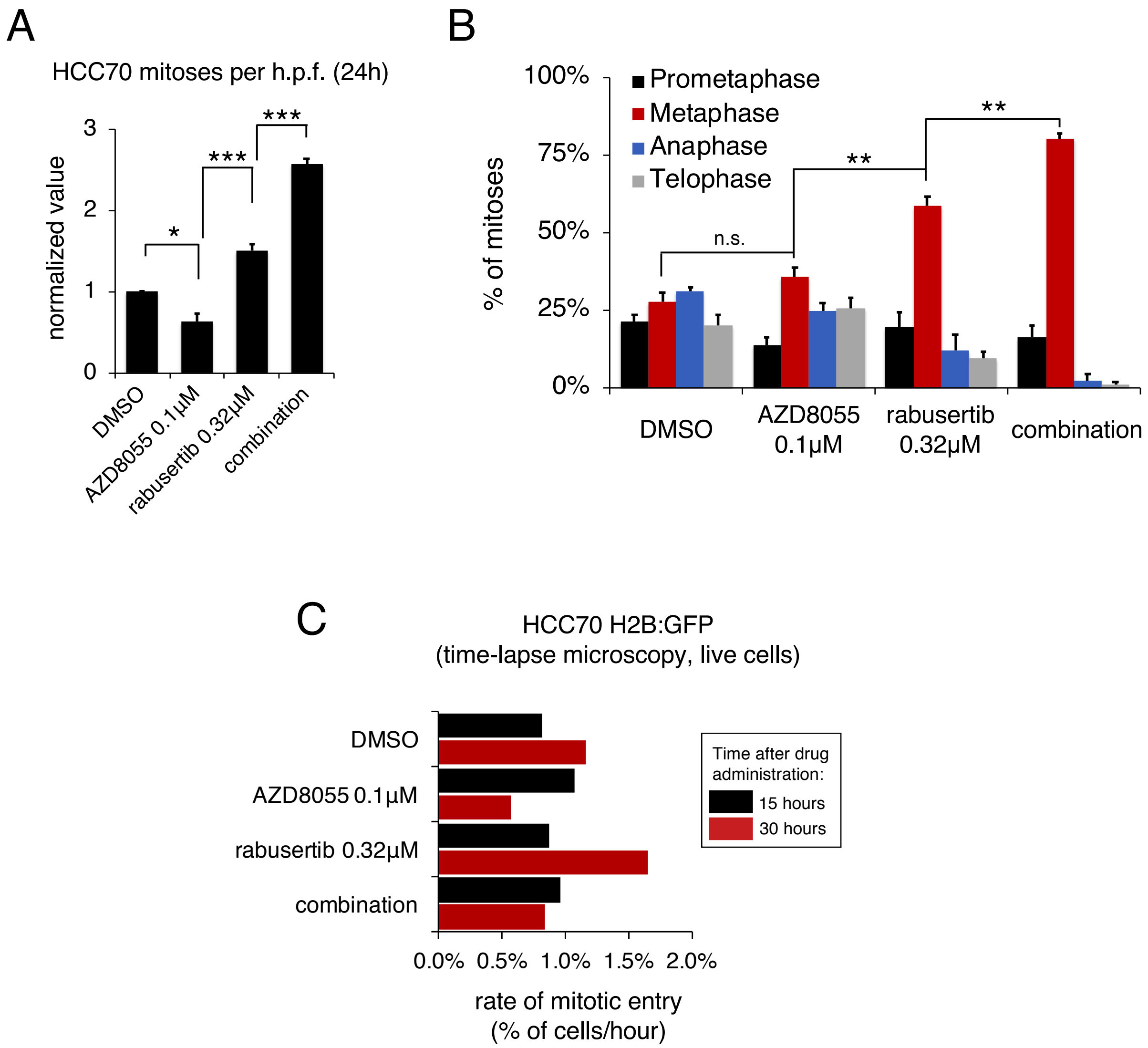
A
HCC70 mitoses per h.p.f. (24h)
normalized value
0
1
2
3
*
***
***
DMSO
AZD8055 0.1µM
rabusertib 0.32µM
combination
B
% of mitoses
0%
25%
50%
75%
100%
Prometaphase
Metaphase
Anaphase
Telophase
n.s.
**
**
DMSO
AZD8055 0.1µM
rabusertib 0.32µM
combination
C
HCC70 H2B:GFP
(time-lapse microscopy, live cells)
DMSO
AZD8055 0.1µM
rabusertib 0.32µM
combination
Time after drug administration:
15 hours
30 hours
0.0%
0.5%
1.0%
1.5%
2.0%
rate of mitotic entry
(% of cells/hour)

**Table S1 (related to Figure 1)**: Classification of breast cancer and non-transformed mammary epithelial cell lines used in this study. 46 cell lines used to assess PTEN and INPP4B protein levels by quantitative western blotting are marked with #. Six TNBC/basal-like cell lines with PI3K pathway activation used for quantitative analysis of responses to PI3K pathway inhibitors appear in **BOLD.** 28 cell lines (*i.e.*, 19 basal-like, 7 luminal and 2 non-transformed mammary epithelial cell lines) used for follow-up studies of drug sensitivity are marked with &.

| Cell lines | Gene expression | *HER2* amplified |
|---|---|---|
| 184-B5[#] | Basal[4] | n/a |
| AU-565[#] | Luminal[1] | Y[1] |
| **BT-20**[#,&] | Basal A[1] | N[1] |
| BT-474[#] | Luminal[1] | Y[1] |
| BT-483[#] | Luminal[1] | N[1] |
| **BT-549**[#,&] | Basal B[1] | N[1] |
| CAL-120[&] | Basal B[5] | n/a |
| CAL-51[&] | Basal B[2] | N[2] |
| CAL-85-1[&] | Basal-like 2[3] | n/a |
| CAMA-1[#,&] | Luminal[1] | N[1] |
| HCC1143[&] | Basal A[1] | N[1] |
| HCC1395[#,&] | Basal B[2] | N[2] |
| HCC1419[#] | Luminal[2] | Y[2] |
| HCC1428[#,&] | Luminal[1] | N[1] |
| HCC1500[#,&] | Luminal[2] | N[1] |
| HCC1569[#] | Basal A[1] | Y[1] |
| **HCC1806**[#,&] | Basal-like 2[3] | N[2] |
| HCC1937[#,&] | Basal A[1] | N[1] |
| HCC1954[#] | Basal A[1] | Y[1] |
| HCC202[#] | Luminal[2] | Y[1] |
| **HCC38**[#,&] | Basal B[1] | N[1] |
| **HCC70**[#,&] | Basal A[1] | N[1] |
| hTERT-HME1[&] | Basal B[2] | N[2] |
| **Hs 578T**[#,&] | Basal B[1] | N[1] |
| MCF-10A[#,&] | Basal B[1] | N[1] |
| MCF10DCIS[#] | n/a | n/a |
| MCF-10F[#] | n/a | n/a |
| MCF-12A[#] | Basal B[1] | N[1] |
| MCF-7[#,&] | Luminal[1] | N[1] |
| MDA-MB-134[#,&] | Luminal[1] | N[1] |
| MDA-MB-157[#,&] | Basal B[1] | N[1] |
| MDA-MB-175[#] | Luminal[1] | N[1] |
| MDA-MB-231[#,&] | Basal B[1] | N[1] |
| MDA-MB-361[#] | Luminal[1] | Y[1] |
| MDA-MB-415[#] | Luminal[1] | N[1] |
| MDA-MB-436[#,&] | Basal B[1] | N[1] |
| MDA-MB-453[#,&] | Luminal[1] | N[1] |
| MDA-MB-468[#,&] | Basal A[1] | N[1] |
| SK-BR-3[#] | Luminal[1] | Y[1] |
| SUM102PT[#] | Basal B[2] | N[2] |
| SUM1315[#,&] | Basal B[1] | N[1] |
| SUM149PT[#,&] | Basal B[1] | N[1] |
| SUM159PT[#,&] | Basal B[1] | N[1] |
| SUM225CWN[#] | Basal A[1] | Y[1] |
| SUM44PE[#] | Luminal[1] | Y[2] |
| SUM52PE[#] | Luminal[1] | Y[1] |
| T-47D[#,&] | Luminal[1] | N[1] |
| UACC-812[#] | Luminal[1] | Y[1] |
| UACC-893[#] | Luminal[2] | Y[1] |
| ZR-75-1[#] | Luminal[1] | N[1] |
| ZR-75-30[#] | Luminal[1] | Y[1] |

**Table S2 (related to Figure 1)**: Molecular characteristics of the six TNBC/basal-like cell lines selected for quantitative analysis of responses to PI3K pathway inhibitors.

| TNBC cell lines | PI3K pathway (mutated gene) | Nucleotide variant | Amino acid substitution | PTEN, INPP4B protein level (rank)[4] | *TP53* mutation[5] (cDNA, protein) |
|---|---|---|---|---|---|
| BT-20 | *PIK3CA* | c.3140A>G[1]<br>c.1616C>G[1] | p.H1047R<br>p.P539R | 2.44 (37)<br>1.83 (23) | c.394A>C[1]<br>p.K132Q |
| BT-549 | *PTEN* | c.823delG[1] | p.V275fs*1 | -1.85 (42)<br>-1.41 (36) | c.747G>C[1]<br>p.R249S |
| HCC1806 | *INPP4B* | c.2541C>G[2] | p.D847E[3] | 2.70 (34)<br>-1.29 (34) | c.766_767insAA[1]<br>p.T256fs*90 |
| HCC38 | *PIK3CA* | c.1157G>T[2] | p.W386L | -1.85 (41)<br>-2.62 (40) | c.818G>T[1]<br>p.R273L |
| HCC70 | *PTEN* | c.270delT[1] | p.F90fs*9 | -2.62 (43)<br>1.74 (25) | c.743G>A[1]<br>p.R248Q |
| Hs 578T | *PIK3R1* | c.1358_1359 insTAA[1] | p.N453_T454 insN | 4.57 (10)<br>-4.89 (46) | c.469G>T[1]<br>p.V157F |

[1]Reported by ATCC.
[2]Reported in COSMIC (searched through canSAR 4.0)
[3]Loss of function substitution according to Lopez SM, et al. *Biochem Biophys Res Commun*, 2013.
[4]Log base 2 of the mean protein level assessed by quantitative western blotting. ( ) = rank in a panel of 48 lysates from 46 breast cancer and non-transformed mammary epithelial cell lines (see **Table S1**), with 1 and 48 assigned to the highest and lowest values, respectively.
[5]All cell lines are homozygous for *TP53* mutations.

**Table S3 (related to Figure 1):** Drug compounds used in this study (N=31). The initial analysis of drug responses used 23 PI3K pathway inhibitors and the MEK inhibitor trametinib (see **BOLD**). Sapanisertib was added during follow-up studies. Six inhibitors of DNA damage response pathways were also used in follow-up studies.

| Compound | Generic name | Clinical development | Phase[#] | Nominal targets |
|---|---|---|---|---|
| **AS-605240** | n/a | N | n/a | PI3K p110γ |
| AZD2281 | olaparib | Y | 1-3;<br>FDA-approved | PARP1/2 |
| **AZD6482** | n/a | Y[1] | 1 | PI3K p110β |
| **AZD8055** | n/a | Y | 1[#] | mTORC1/2 |
| AZ20 | n/a | N | n/a | ATR |
| **BEZ235** | dactolisib | Y | 1-2 | PI3K p110(α,β,δ,γ), mTORC1/2, ATR |
| **BKM120** | buparlisib | Y | 1-3 | PI3K p110(α,β,δ,γ) |
| **BYL719** | alpelisib | Y | 1-3 | PI3K p110α |
| **BX-912** | n/a | N | n/a | PDK1 |
| **CAL-101**<br>(GS-1101) | idelalisib | Y | 1-3;<br>FDA-approved | PI3K p110δ |
| **GDC-0941** | pictilisib | Y | 1-2[#] | PI3K p110(α,β,δ,γ) |
| **GDC-0980** | apitolisib | Y | 1-2 | PI3K p110(α,β,δ,γ), mTORC1/2 |
| **GSK1120212** | trametinib | Y | 1-3;<br>FDA-approved | MEK1/2 |
| **GSK2126458** | omipalisib | Y | 1[#] | PI3K p110(α,β,δ,γ), mTORC1/2 |
| **KIN001-244** | n/a | N | n/a | PDK1 |
| KU-60019 | n/a | N | n/a | ATM |
| LY2603618 | rabusertib | Y | 1-2[#] | Chk1 |
| **MK-2206** | n/a | Y | 1-2 | Akt1/2/3 |
| MLN0128<br>(INK-128) | sapanisertib | Y | 1-2 | mTORC1/2 |
| NU7026 | n/a | N | n/a | DNA-PK |
| **PI-103** | n/a | N | n/a | PI3K p110(α,β,δ,γ), mTORC1/2, DNA-PK |
| **PIK-93** | n/a | N | n/a | PI3K p110α, p110γ; PI4KIIIβ |
| **RAD001** | everolimus | Y | 1-3;<br>FDA-approved | mTORC1 |
| **rapamycin** | sirolimus | Y | 1-3 | mTORC1 |
| **TGX-221** | n/a | N | n/a | PI3K p110β |
| **Torin1** | n/a | N | n/a | mTORC1/2, DNA-PK |
| **Torin2** | n/a | N | n/a | mTORC1/2, ATR, ATM, DNA-PK |
| VE-821 | n/a | N | n/a | ATR |
| **WYE-132** | n/a | N | n/a | mTORC1/2 |
| **XL147**<br>(SAR245408) | pilaralisib | Y | 1-2 | PI3K p110(α,β,δ,γ) |
| **ZSTK474** | n/a | Y | 1[#] | PI3K p110(α,β,δ,γ) |

[#]Indicates no active clinical trials (clinicaltrials.gov).
[1]Studied in patients for anti-platelet effects.

**Table S4 (related to Figure 2):** Downregulated (blue) and upregulated (red) metabolites in BT-549 cells following 6 hours of exposure to 1μM rapamycin, AZD8055, or Torin2, as identified by targeted metabolomics profiling. Analysis was performed using MetaboAnalyst 4.0 (see STAR methods for details).

**Rapamycin** (volcano plot cut-offs: fold change 1.5 and p-value 0.12):

| metabolite | KEGG/HMDB ID | metabolic pathway | fold change | $log_2$ (fold change) | raw p value | $-log_{10}$ (p value) |
|---|---|---|---|---|---|---|
| 1,3-diphosphateglycerate | C00236 | glycolysis | 0.17506 | -2.5141 | 0.005024 | 2.2989 |
| glycerophosphocholine | C00670 | glycerophospholipid metabolism | 0.30941 | -1.6924 | 0.006531 | 2.185 |
| 2,3-diphosphoglyceric acid | C01159 | glycolysis | 0.42366 | -1.239 | 0.088396 | 1.0536 |
| UTP-nega | C00075 | pyrimidine metabolism | 0.47819 | -1.0644 | 0.079647 | 1.0988 |
| dTDP-nega | C00363 | pyrimidine metabolism | 0.49717 | -1.0082 | 0.1124 | 0.94924 |
| dGTP | C00286 | purine metabolism | 0.51058 | -0.96978 | 0.096791 | 1.0142 |
| NADPH-nega | C00005 | glutathione metabolism | 0.5184 | -0.94785 | 0.057813 | 1.238 |
| CDP-choline | C00307 | glycerophospholipid metabolism | 0.56569 | -0.82192 | 0.057738 | 1.2385 |
| CMP | C00055 | pyrimidine metabolism | 0.57775 | -0.79148 | 0.10558 | 0.97642 |
| carbamoyl phosphate | C00169 | pyrimidine metabolism | 0.59182 | -0.75676 | 0.10358 | 0.98473 |
| 5-methoxytryptophan | HMDB02339 | tryptophan metabolism | 0.60642 | -0.72162 | 0.034779 | 1.4587 |
| glutamine | C00064 | purine/pyrimdine metabolism, others | 1.5486 | 0.63099 | 0.037744 | 1.4232 |
| p-hydroxybenzoate | C00156 | benzoate degradation | 1.6104 | 0.68743 | 0.078678 | 1.1041 |
| kynurenine | C00328 | tryptophan metabolism | 1.7086 | 0.77283 | 0.081955 | 1.0864 |
| cytosine | C00380 | pyrimidine metabolism | 1.8465 | 0.8848 | 0.012569 | 1.9007 |
| N-acetylputrescine | C02714 | arginine and proline metabolism | 2.004 | 1.0029 | 0.028281 | 1.5485 |
| cytidine | C00475 | pyrimidine metabolism | 2.0479 | 1.0341 | 0.013472 | 1.8706 |
| dTMP | C00364 | pyrimidine metabolism | 2.5661 | 1.3596 | 0.031743 | 1.4983 |

Metabolites detectable in all DMSO control samples but not in all treated samples (shown are peak intensity values):

| metabolites | DMSO_1 | DMSO_2 | DMSO_3 | Rapamycin_1 | Rapamycin_2 | Rapamycin_3 |
|---|---|---|---|---|---|---|
| FMN | 13817.8342 | 9204.51331 | 13128.728 | 6150.84896 | NA | NA |
| cyclic bis(3->5) dimeric GMP | 7676.66828 | 3390.80843 | 14429.9837 | NA | NA | NA |
| succinyl-CoA-methylmalonyl-CoA-nega | 5090.87455 | 5085.01944 | 8670.18606 | NA | NA | NA |
| diiodothyronine | 8350.47985 | 6784.31051 | 4675.54214 | NA | NA | NA |

**AZD8055** (volcano plot cut-offs: fold change 1.5 and p-value 0.12):

| metabolite | KEGG/HMDB ID | metabolic pathway | fold change | $log_2$ (fold change) | raw p value | $-log_{10}$ (p value) |
|---|---|---|---|---|---|---|
| 2,3-diphosphoglyceric acid | C01159 | glycolysis | 0.33203 | -1.5906 | 0.052983 | 1.2759 |
| N-carbamoyl-L-aspartate-nega | C00438 | pyrimidine metabolism | 0.57796 | -0.79095 | 0.11241 | 0.94921 |
| dGDP-nega | C00361 | purine metabolism | 0.6014 | -0.73361 | 0.11961 | 0.92221 |
| dTDP-nega | C00363 | pyrimidine metabolism | 0.61724 | -0.6961 | 0.083038 | 1.0807 |
| N-acetyl-L-alanine | C01073 | beta-alanine metabolism | 0.65304 | -0.61475 | 0.11847 | 0.92639 |
| 4-aminobutyrate | C00334 | alanine, aspartate & glutamate metabolism | 0.65956 | -0.60043 | 0.054075 | 1.267 |
| IDP-nega | C00104 | purine metabolism | 0.66408 | -0.59057 | 0.11967 | 0.92201 |
| phenylpropiolic acid | HMDB00563 | phenylalanine metabolism | 1.6197 | 0.69575 | 0.10052 | 0.99776 |
| 3-S-methylthiopropionate | C08276 | cysteine & methionine metabolism | 1.6749 | 0.74411 | 0.0093521 | 2.0291 |
| 5-methyl-THF | C00440 | one carbon pool by folate | 1.7904 | 0.8403 | 0.11602 | 0.93546 |
| malonyl-CoA-posi | C00083 | fatty acid biosynthesis | 1.8578 | 0.89362 | 0.017418 | 1.759 |
| N-acetylputrescine | C02714 | arginine & proline metabolism | 2.0599 | 1.0426 | 0.071022 | 1.1486 |
| dTMP | C00364 | pyrimidine metabolism | 2.0744 | 1.0527 | 0.044968 | 1.3471 |
| cytosine | C00380 | pyrimidine metabolism | 2.1107 | 1.0778 | 0.0002731 | 3.5638 |
| deoxyinosine | C05512 | purine metabolism | 2.133 | 1.0929 | 0.020202 | 1.6946 |
| D-sedoheptulose-1-7-phosphate | C05382 | pentose phosphate pathway | 2.7239 | 1.4457 | 0.022382 | 1.6501 |
| cytidine | C00475 | pyrimidine metabolism | 3.0225 | 1.5958 | 0.0001058 | 3.9754 |
| S-methyl-5-thioadenosine | C00170 | cysteine & methionine metabolism | 3.2664 | 1.7077 | 0.001777 | 2.7503 |
| hypoxanthine | C00262 | purine metabolism | 4.9622 | 2.311 | 0.05315 | 1.2745 |

Metabolites detectable in all DMSO control samples but not in all treated samples (shown are peak intensity values):

| metabolites | DMSO_1 | DMSO_2 | DMSO_3 | AZD8055_1 | AZD8055_2 | AZD8055_3 |
|---|---|---|---|---|---|---|
| FMN | 13817.834 | 9204.5133 | 13128.728 | 12843.701 | NA | NA |
| cyclic bis(3->5) dimeric GMP | 7676.6683 | 3390.8084 | 14429.984 | NA | NA | NA |
| succinyl-CoA-methylmalonyl-CoA-nega | 5090.8746 | 5085.0194 | 8670.1861 | NA | NA | NA |
| glycine | 26286.235 | 7645.5868 | 19907.092 | NA | NA | 16124.418 |
| choline | 2871318.1 | 804003.85 | 2266906.1 | 4928666.6 | NA | NA |
| 5-methoxytryptophan | 11870.832 | 7628.6091 | 13568.743 | 5948.8763 | NA | NA |
| diiodothyronine | 8350.4798 | 6784.3105 | 4675.5421 | NA | NA | NA |
| retinoic acid | 14769.437 | 6658.804 | 3700.0403 | NA | NA | 3813.5295 |

#### Table S4 (related to Figure 2), *continued*

**Torin2** (volcano plot cut-offs: fold change 1.5 and p-value 0.12):

| metabolite | KEGG/HMDB ID | metabolic pathway | fold change | $log_2$ (fold change) | raw p value | $-log_{10}$ (p value) |
|---|---|---|---|---|---|---|
| NADH | C00004 | oxidative phosphorylation | 0.30588 | -1.709 | 0.094691 | 1.0237 |
| dTDP-nega | C00363 | pyrimidine metabolism | 0.31052 | -1.6873 | 0.0073083 | 2.1362 |
| glyoxylate | C00048 | glyoxylate & dicarboxylate metabolism | 0.31628 | -1.6607 | 0.066986 | 1.174 |
| nicotinamide ribotide | C00455 | nicotinate & nicotinamide metabolism | 0.42276 | -1.2421 | 0.010448 | 1.981 |
| dCTP-nega | C00458 | pyrimidine metabolism | 0.46269 | -1.1119 | 0.004728 | 2.3253 |
| dGTP | C00286 | purine metabolism | 0.51939 | -0.9451 | 0.099317 | 1.003 |
| dGDP-nega | C00361 | purine metabolism | 0.53069 | -0.91406 | 0.077495 | 1.1107 |
| ADP-nega | C00008 | purine metabolism | 0.55566 | -0.84773 | 0.094017 | 1.0268 |
| NADPH-nega | C00005 | glutathione metabolism | 0.61037 | -0.71225 | 0.099435 | 1.0025 |
| IDP-nega | C00104 | purine metabolism | 0.63205 | -0.66189 | 0.095918 | 1.0181 |
| aminoimidazole carboxamide ribonucleotide | C04677 | purine metabolism | 0.64544 | -0.63164 | 0.021033 | 1.6771 |
| atrolactic acid | HMDB00475 | phenylalanine metabolism | 1.5241 | 0.60794 | 0.075853 | 1.12 |
| thymine | C00178 | pyrimidine metabolism | 1.573 | 0.6535 | 0.018955 | 1.7223 |
| D-glucosamine-6-phosphate | C00352 | amino sugar & nucleotide sugar metabolism | 1.6124 | 0.68925 | 0.11918 | 0.92381 |
| nicotinamide riboside | C03150 | nicotinate & nicotinamide metabolism | 1.6263 | 0.70158 | 0.0008533 | 3.0689 |
| cytosine | C00380 | pyrimidine metabolism | 1.9227 | 0.94312 | 0.0029985 | 2.5231 |
| deoxyguanosine | C00330 | purine metabolism | 1.9346 | 0.95202 | 0.10065 | 0.99717 |
| N-acetylputrescine | C02714 | arginine & proline metabolism | 2.0606 | 1.0431 | 0.052397 | 1.2807 |
| cytidine | C00475 | pyrimidine metabolism | 2.2522 | 1.1714 | 0.0002413 | 3.6175 |
| putrescine | C00134 | arginine & proline metabolism | 2.2827 | 1.1908 | 0.0058995 | 2.2292 |
| D-sedoheptulose-1-7-phosphate | C05382 | pentose phosphate pathway | 2.4255 | 1.2783 | 0.087554 | 1.0577 |
| dTMP | C00364 | pyrimidine metabolism | 3.5606 | 1.8321 | 0.0048334 | 2.3157 |
| hypoxanthine | C00262 | purine metabolism | 5.5227 | 2.4654 | 0.044853 | 1.3482 |

Metabolites detectable in all DMSO control samples but not in all treated samples (shown are peak intensity values):

| metabolites | DMSO_1 | DMSO_2 | DMSO_3 | Torin2_1 | Torin2_2 | Torin2_3 |
|---|---|---|---|---|---|---|
| FMN | 13817.834 | 9204.5133 | 13128.728 | NA | NA | 5086.4902 |
| cyclic bis(3->5) dimeric GMP | 7676.6683 | 3390.8084 | 14429.984 | 4185.0821 | NA | NA |
| succinyl-CoA-methylmalonyl-CoA-nega | 5090.8746 | 5085.0194 | 8670.1861 | NA | NA | NA |
| choline | 2871318.1 | 804003.85 | 2266906.1 | NA | NA | NA |
| histidinol | 15249.437 | 5476.1406 | 10984.721 | NA | NA | 9324.3814 |
| flavone | 8964.2054 | 7859.0649 | 7875.8126 | NA | NA | NA |
| 5-methoxytryptophan | 11870.832 | 7628.6091 | 13568.743 | NA | NA | NA |
| diiodothyronine | 8350.4798 | 6784.3105 | 4675.5421 | 16973.513 | NA | NA |
| retinoic acid | 14769.437 | 6658.804 | 3700.0403 | NA | NA | 8282.8993 |

**Summary and interpretation**: Compared to BT-549 cells treated with DMSO, cells treated with active drug for 6h exhibit decreased levels of both pyrimidine and purine deoxyribonucleotide triphosphates and/or their precursors including deoxycytidine triphosphate (dCTP), cytidine monophosphate (CMP), deoxythymidine diphosphate (dTDP), deoxyguanosine triphosphate (dGTP) and adenosine diphosphate (ADP), among others. The changes in metabolite levels are consistent with inhibition of both *de novo* synthesis and salvage pathways for pyrimidine and purine nucleotides.

- Increased levels of cytidine and cytosine suggest inhibition of pyrimidine salvage pathways.
- Decreased levels of carbamoyl phosphate, N-carbamoyl-L-aspartate and/or uridine triphosphate (UTP) suggest decreased *de novo* synthesis of pyrimidines.
- Increased levels of deoxythymidine monophosphate (dTMP) and decreased levels of deoxythymidine diphosphate (dTDP) are consistent with reduced activity of deoxythymidylate kinase, an enzyme that acts downstream of *de novo* synthesis and salvage pathways for pyrimidines.
- Increased levels of hypoxanthine, deoxyinosine and/or deoxyguanosine suggest inhibition of purine salvage pathways.
- Decreased levels of AICAR may reflect decreased *de novo* purine synthesis.
